# Functional convergence of autonomic and sensorimotor processing in the lateral cerebellum

**DOI:** 10.1101/683573

**Authors:** Vincenzo Romano, Aoibhinn L. Reddington, Silvia Cazzanelli, Mario Negrello, Laurens W.J. Bosman, Chris I. De Zeeuw

**Affiliations:** Department of Neuroscience, Erasmus MC, Rotterdam, The Netherlands; Netherlands Institute for Neuroscience, Royal Academy of Arts and Sciences, Amsterdam, The Netherlands

## Abstract

The cerebellum is involved in control of voluntary and autonomic rhythmic behaviors, yet it is largely unclear to what extent it coordinates these in a concerted action. Here, we studied Purkinje cell activity during unperturbed and perturbed respiration in cerebellar lobules simplex, crus 1 and 2. During unperturbed (eupneic) respiration complex spike and simple spike activity encoded respiratory activity, the timing of which corresponded with ongoing sensorimotor feedback. Instead, upon whisker stimulation mice concomitantly accelerated their simple spike activity and inspiration in a phase-dependent manner. Moreover, the accelerating impact of whisker stimulation on respiration could be mimicked by optogenetic stimulation of Purkinje cells and prevented by cell-specific genetic modification of their AMPA receptors that hampered increases in simple spike firing. Thus, the impact of Purkinje cell activity on respiratory control is context- and phase-dependent, suggesting a coordinating role for the cerebellar hemispheres in aligning autonomic and sensorimotor behaviors.

## Introduction

Rhythmic behaviors are part of everyday life of mammals. They can emerge from predominantly conscious activity such as locomotion, licking or whisking, but also from more subconscious behaviors like heart beat or respiration. Speed, amplitude and phase of rhythmic movements depend on the behavioral demands and context, and thereby they depend on each other (Welker, 1964; Cao et al., 2012; Moore et al., 2013; Kurnikova et al., 2017). Accordingly, many of the motor domains involved in rhythmic movements serve multiple functions and many of these functions can be coordinated in a concerted action. For example, inspiration is driven by the diaphragm and intercostal muscles, which are also involved in postural control (Rimmer et al., 1995; Hodges and Gandevia, 2000), and respiration and posture are synergistically controlled during processes like vocalization, swimming or parturition (Tomori and Widdicombe, 1969; Jakovljevic and McConnell, 2009; Holstege, 2014).

When different forms of sensorimotor behaviors have to be coordinated, the olivocerebellar system is often involved in optimal fine-tuning in time and space (Kitazawa et al., 1998; Vinueza Veloz et al., 2015; Owens et al., 2018). This presumably not only holds for non-rhythmic behaviors, but also for rhythmic behaviors like respiration (Gozal et al., 1995; Parsons et al., 2001; Isaev et al., 2002; McKay et al., 2003; Cao et al., 2012; Raux et al., 2013; Critchley et al., 2015; Park et al., 2016). Accordingly, rare, but dramatic cases of sudden infant death syndrome (SIDS) have been attributed to acute respiratory arrest in relation to inferior olivary hypoplasia or delayed maturation of the cerebellar cortex (Cortez and Kinney, 1996; Cruz-Sánchez et al., 1997; Harper, 2000; Lavezzi et al., 2013; Katsetos et al., 2014), while cerebellar dysfunction has been observed in congenital central hypoventilation syndrome, which entails the inability to react to dyspnea (Harper et al., 2005; Kumar et al., 2008; Harper et al., 2015). Likewise, patients with a cerebellar tumor or hemorrhage frequently need mechanical ventilation, often showing a relatively slow recovery of respiration after neurosurgery (Chen et al., 2005; Tsitsopoulos et al., 2012; Lee et al., 2013; Abecassis et al., 2017; Arnone et al., 2017). Moreover, most cerebellar ataxia patients have trouble to modulate their breathing during exercise (Ebert et al., 1995; Deger et al., 1999; De Joanna et al., 2008). Thus, there is ample evidence for a role of the olivocerebellar system in controlling respiration and adjusting it according to behavioral demands, pointing towards a role in synergistic integration of autonomic and voluntary behaviors.

At present, it is unclear to what extent different rhythmic behaviors can be controlled by the same cerebellar region and cells, and if so, how they might contribute to synergistic control of the different motor domains involved. Here, we studied the activity of Purkinje cells in the lateral cerebellum in relation to respiratory control, while interfering with their whisker system. We focused on lobule simplex in conjunction with lobules crus 1 and crus 2, because they are strongly related to rhythmic whisker movements and because their cells have been shown to respond to a variety of somatosensory inputs from the face, possibly integrating different sensorimotor behaviors (Shambes et al., 1978; Bosman et al., 2010; Chen et al., 2016; Brown and Raman, 2018; Romano et al., 2018; Ju et al., 2019). We found Purkinje cells in these lobules co-modulate their firing rate with multiple phases of the respiratory cycle during unperturbed breathing. When we briefly stimulated the whiskers, the mice advanced the phase of their simple spike activity and breathing behavior concomitantly. The Purkinje cells that responded to whisker stimulation and also contributed to acceleration of respiration were particularly prominent in medial crus 1. Moreover, the respiratory adjustment following whisker stimulation could be induced by transiently stimulating these Purkinje cells in the lateral cerebellum optogenetically, whereas it was significantly impaired following Purkinje cell-specific impairment of postsynaptic AMPA receptors. Together, our data implicate that the cerebellar hemispheres can control respiratory behavior and align its rhythm with that of other behaviors in a phase-dependent manner, highlighting their putative role in synergistic integration of different sensorimotor activities.

## Results

### Unperturbed respiratory behavior

To find out to what extent Purkinje cells in the cerebellar hemispheres encode the three phases of unperturbed (eupneic) respiration, defined as a cycle of inspiration, post-inspiration and expiration (Richter and Smith, 2014; Anderson and Ramirez, 2017), we studied their activity patterns in awake head-restrained mice. During inspiration, contractions of the diaphragm and external intercostal muscles generate a volume expansion of the lungs, while during post-inspiration the inspiration muscles relax and laryngeal constriction muscles retard lung compression (Dutschmann and Paton, 2002). During active expiration, abdominal and internal intercostal muscles contract depending upon metabolic demand (Aliverti et al., 1997; Bianchi and Gestreau, 2009). The respiration phases of awake head-restrained mice were measured with a pressure sensor placed under the abdomen and analyzed upon phase transformation. Under these conditions, the mice had a median breathing frequency of 2.4 Hz (inter-quartile range (IQR): 1.0 Hz, n = 13 mice), with a median CV of 0.51 (IQR: 0.35), indicating the fast nature as well as substantial level of variability of their breathing rhythm at rest (Fig. S1a-c).

### Purkinje cell complex spike activity during unperturbed respiration

In the first set of experiments we recorded the activity of 43 Purkinje cells during unperturbed respiration. These cells had a median complex spike firing frequency of 1.37 Hz (IQR: 0.54 Hz) (Fig. S1d). The extent of complex spike firing rate modulation along the respiratory cycle was quantified per Purkinje cell by comparing the measured distributions of complex spikes with randomly shuffled ones. The random shuffling was performed 500 times, upon which the 99% confidence interval (Z = 3) was calculated. Firing patterns exceeding this 99% confidence interval were considered indicative of a statistically significant modulation (Fig. 1a, Fig. S1e). Of the 43 Purkinje cells, 9 (21%) displayed a statistically significant complex spike modulation, but also many of the other Purkinje cells showed some degree of modulation (Fig. S1e). Of all 43 Purkinje cells, 19 (44%) displayed maximum complex spike firing in the period just before 3π/2 (Fig. 1b), which is around the transition from post-inspiration to expiration, while the other Purkinje cells typically peaked at a given phase during inspiration or post-inspiration, but never during expiration.

**Figure 1.**
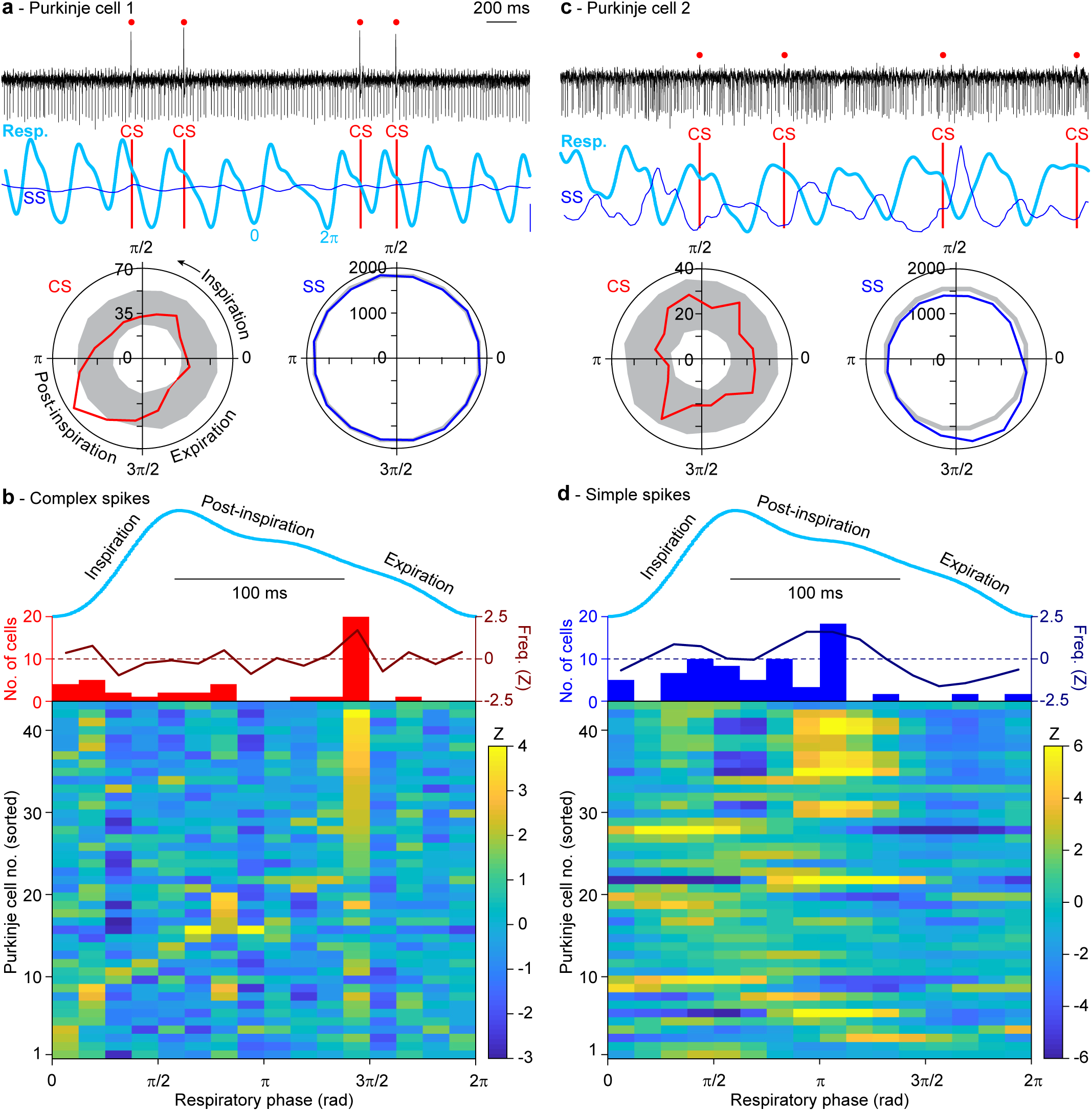
Purkinje cells in the lateral cerebellum encode eupneic breathing. **a** An example recording of a Purkinje cell showing complex spike (CS), but no simple spike (SS), modulation during unperturbed (eupneic) respiration in an awake mouse. The complex spikes are indicated by red dots and vertical lines. The instantaneous simple spike rate (thin blue line) is indicated in combination with the respiratory signal (thick cyan line). The blue scale bar on the right indicates 25 Hz of simple spike modulation. The raw signal indicates that the complex spikes preferably occurred around the transition from post-inspiration to expiration as confirmed by a polar plot summarizing the whole recording. The gray areas indicate the 99% confidence interval after bootstrap. **b** Of the 43 Purkinje cells recorded during eupneic respiration, 19 displayed their maximal complex spike firing around 3π**/**2. This is illustrated as the average modulation in firing rate (red line, middle), the distribution of the phases of strongest modulations (histogram; middle) and a heat map illustrating the complex spike firing patterns of 43 Purkinje cells (bottom). This analysis was performed without pre-selection of Purkinje cells. For comparison, a randomly chosen respiratory cycle is indicated (cyan). Note that the respiratory trace is plotted based on time, while the heat map and histogram are based on the phase. **c** An example of another Purkinje cell, showing relatively weak complex spike modulation, but strong simple spike modulation during eupneic respiration. **d** The same analysis as in **b**, but for the simple spikes, revealing a preference for simple spike firing during post-inspiration (just before the complex spike peak) and a relatively low firing rate during expiration (following the complex spike peak). Note that the Purkinje cells of both heat maps are sorted by the phase of the maximal increase in complex spike firing. Consequently, the cell numbers of **b** and **d** refer to the same Purkinje cells.

### Purkinje cell simple spike activity during unperturbed respiration

The median simple spike rate of the 43 Purkinje cells was 64.54 (IQR: 34.93) Hz (Fig. S1d). The simple spike activity of the majority of these cells (i.e., 35 or 81%) showed a statistically significant modulation across different phases of respiration (Fig. 1c; Fig. S1e). Compared to the modulation of complex spike firing, the preferred phases of the peaks of the simple spike modulation were more closely associated with the inspiration and early post-inspiration periods (Fig. 1d). When we considered the absolute timing – rather than the phase – of the modulation of simple spike activity of single Purkinje cells relative to the start of inspiration, we found that the simple spike rate modulation of 26 Purkinje cells exceeded a Z criterion of higher than 2 (*p* < 0.05). In most of these Purkinje cells, simple spike modulation was essentially bi-directional (Fig. S2a). However, in half of the cells the amplitude of the increase in simple spike modulation was stronger than the decrease, whereas in the other half it was opposite (Fig. 2a-c; Fig. S2a). The peaks of the decreased firing generally preceded those of the increased firing, yielding a population average of a short-latency decrease followed by an increase of simple spike activity (Fig. 2c-e). This order of events of simple spike decreases and increases was substantiated by a positive correlation between the amplitude of the strongest correlation and its time of occurrence (r = 0.49, *p* = 0.010, n = 26, Pearson correlation; Fig. 2d). The preference for decreased or increased firing did not depend on the baseline frequency (r = 0.21, *p* = 0.309, Pearson correlation; Fig. S2b). The population average of simple spike activity approximated the actual respiratory behavior rather well with zero phase-lag, suggesting the relevance of a population encoding mechanism (Fig. 2e).

**Figure 2.**
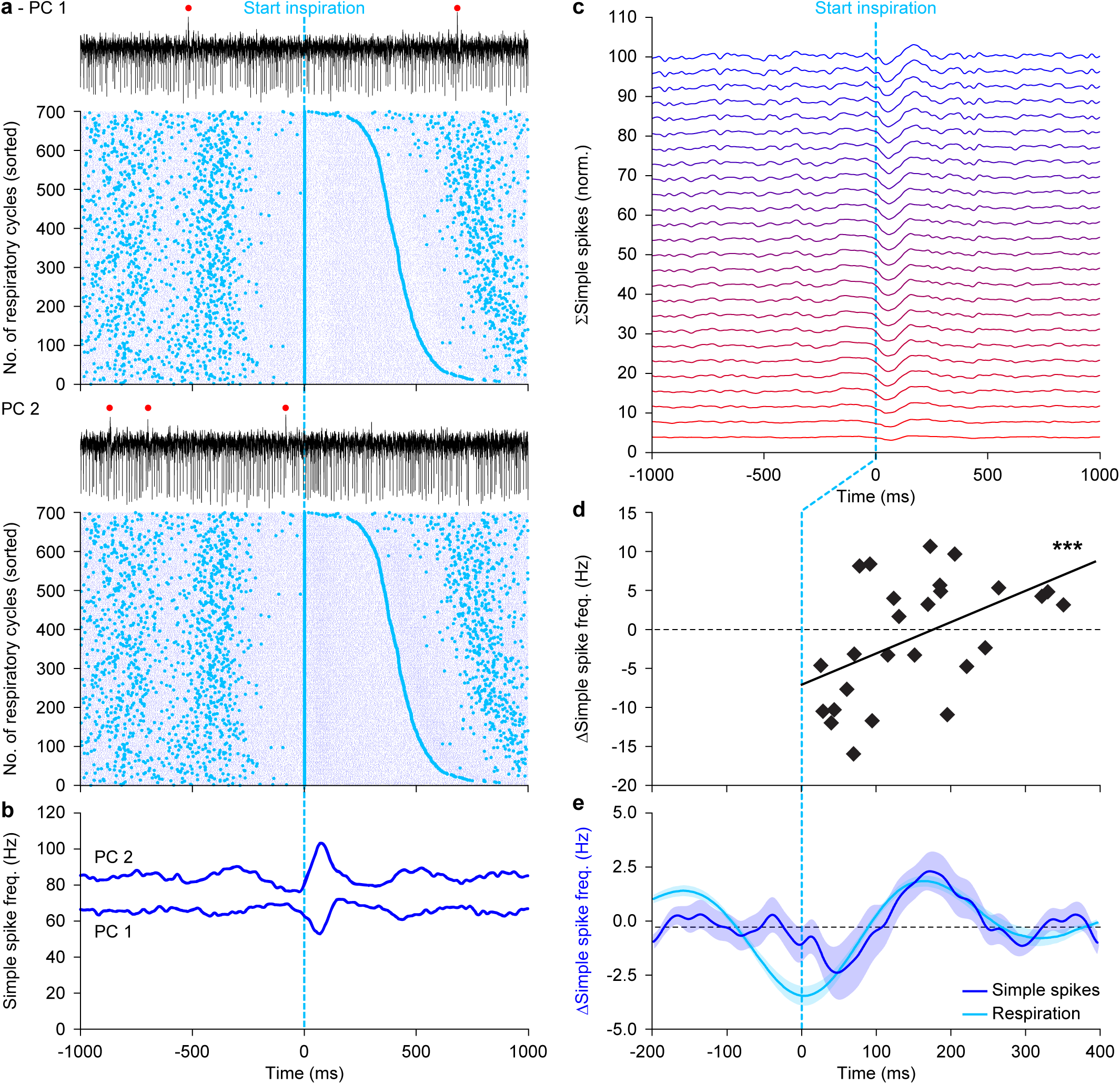
Eupneic respiration is associated with both increased and decreased simple spike firing. **a** During the respiratory cycle, simple spike modulation can either be predominantly decreasing or increasing. This is illustrated by two representative Purkinje cells (PCs) recorded simultaneously in the same animal. The trials in the raster plots are aligned on the start of an inspiration and are sorted according to the duration of the interval to the start of the next inspration. Cyan dots indicate the start of inspiration and blue dots the occurrence of simple spikes. The red dots on top of the traces indicate complex spikes. **b** Convolved peri-stimulus time histograms of the two Purkinje cells shown in **a**. **c** Stacked line plot of the instantaneous simple spike firing rates of all 26 Purkinje cells that displayed a statistically significant modulation of their simple spike firing during unperturbed (eupneic) breathing. The simple spike firing is displayed in percentage of baseline firing and normalized so that the upper line reflects the population average. The Purkinje cells are ordered from the cell with the strongest decrease (bottom, red line) to the strongest increase (top, blue line) in simple spike modulation. Each trace is aligned to the start of inspiraton. **d** Scatter plot of moments of maximal modulation per Purkinje cell, showing a correlation between the time of maximum modulation and its amplitude (r = 0.54, *p* < 0.001, Spearman rank test). Note that on average, as well as at individual cell level, the suppression of simple spikes preceded the increase. **e** Overall, the simple spikes were found to follow, rather than to lead, the respiratory signal. Lines indicate the averages and the shaded areas the SEM.

We further explored whether trial-by-trial variations in simple spike firing correlated with variations in the respiratory signal. We designed a matrix of correlation in which, for each respiratory cycle, the respiratory signal was compared to the instantaneous simple spike rate aligned to the start of each inspiration. This analysis reveals the temporal relationships between both signals whereby a correlation along the 45° line indicates a synchronous event. Indeed, Purkinje cells with a preference for increased simple spike firing as well as those that predominantly show decreased simple spike firing displayed correlations between simple spikes and respiration. The strongest effects were found with the respiratory signal leading the simple spike firing (Fig. 3). Thus, while simple spikes generally co-modulate with the phase of the respiratory signal with approximately a zero lag (Fig. 2e), the depth of their simple spike modulation reflected the depth of the respiration with a delay.

**Figure 3.**
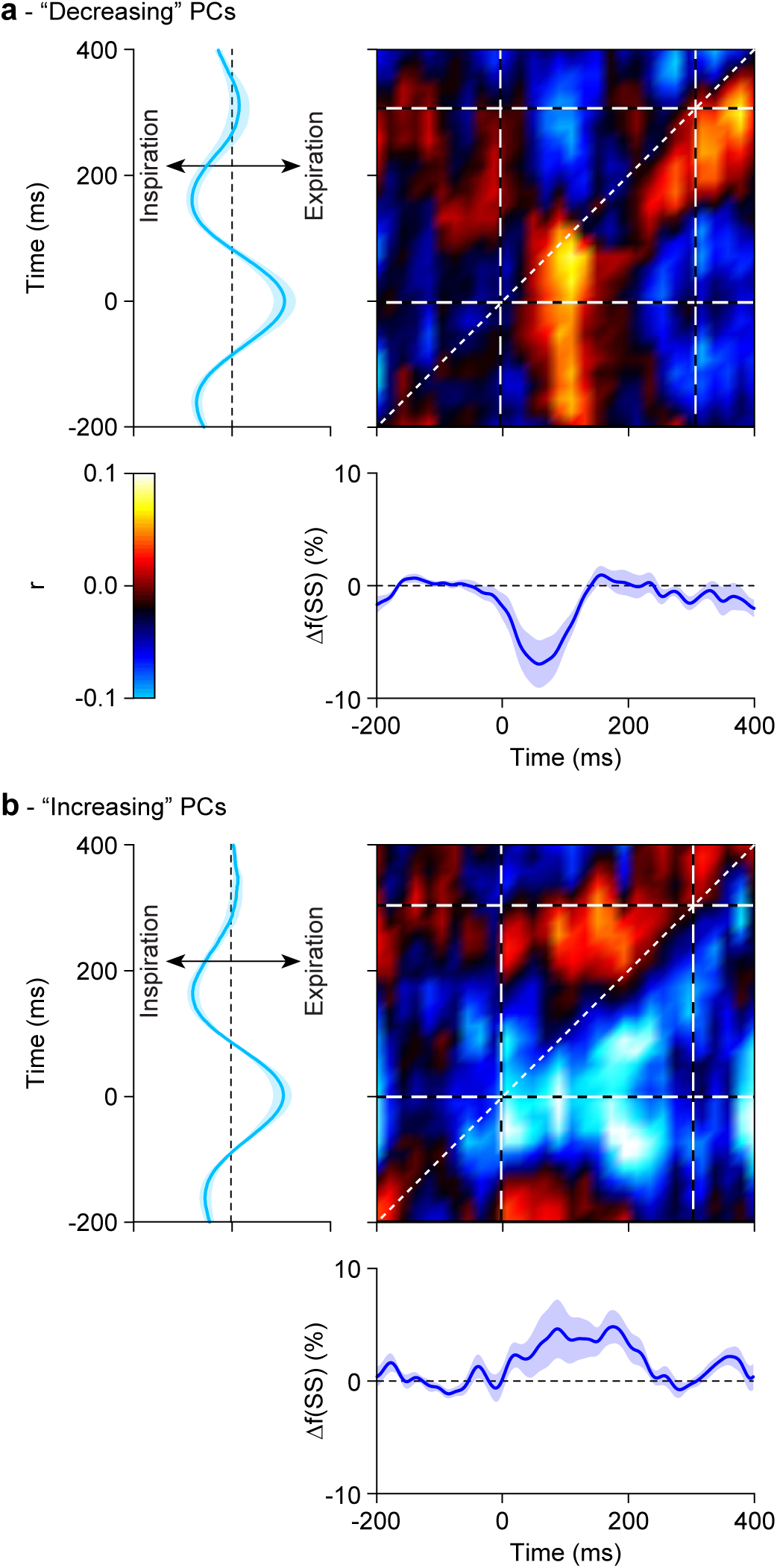
Simple spike modulation follows inspiration. **a** Correlation matrix showing a positive correlation between simple spike firing (blue trace at the bottom shows convolved peri-stimulus time histogram triggered on start of inspiration) and respiration (cyan trace at the left; indicated is the mean ± SEM) based on a trial-by-trial variance analysis in those Purkinje cells that predominantly showed a decrease in their simple spike firing rate linked to the respiratory cycle. Average of 13 Purkinje cells during unperturbed breathing. Note that the main simple spike activity follows the respiration (red area is below the 45° line). **b** Similar analysis of the 13 Purkinje cells that predominantly showed increased simple spike firing during the respiratory cycle. These Purkinje cells had a mostly negative correlation. Thus, for both types of Purkinje cells, the correlation was opposite to their mode of modulation, indicating that the shallower the respiration, the stronger the simple spike modulation.

The complex spike and simple spike modulation of each Purkinje cell typically occurred during different phases of the respiratory cycle, although often not in exact anti-phase (Fig. S3a). The occurrence of increased simple spike firing around π correlated well with increased complex spike firing around 3π/2 (r = 0.536, *p* < 0.001, Spearman rank test; Fig. S3b). In turn, this latter peak in complex spike firing correlated with a subsequent decrease in simple spike firing during expiration (r = −0.431, *p* = 0.004, Spearman rank test; Fig. S3c). Thus, there were signs of reciprocity between complex spike and simple spike firing with a temporal shift of about 50-80 ms, which is reminiscent of studies of other cerebellar regions (Badura et al., 2013; Chaumont et al., 2013; Witter et al., 2013).

### Respiratory behavior following whisker stimulation

Given the intricate relationships between oro-facial behaviors in general and the harmonization of respiratory and whisking behavior in rodents in particular (Lu et al., 2013; Moore et al., 2013; Kurnikova et al., 2017), we wondered how an air puff to the facial whiskers that triggers reflexive whisker protraction (Bellavance et al., 2017; Brown and Raman, 2018; Romano et al., 2018) would also affect the respiratory cycle. To evaluate this, we subjected 12 mice to periodic 0.5 Hz whisker stimulation while measuring their respiration (Fig. 4a-d). When delivered within 100 ms after the start of the previous inspiration, the air puff had little effect, but otherwise it accelerated the start of the next inspiration with a median latency of 91 (IQR: 106) ms (Fig. 4e; Fig. S4a). Thus, stimulation of the whiskers induced a phase-dependent accelerating respiratory response, shortening the interval between the air puff and the start of the next inspiration (*p* < 0.001 for experimental data vs. *p* = 0.719 for randomized data, Kolmogorov-Smirnov test; Fig. 4f-g; Fig. S4a). Rather than entraining their respiratory rhythm to the (fixed) frequency of air puff stimulation, the mice adjusted the timing of inspiration of the respiratory cycle directly following sensory stimulation (Fig. 4h; Fig. S4b). Variations in the level of sensory-induced whisker protraction and depth of respiration were correlated; trial-by-trial variations revealed that stronger whisker protractions preceded deeper levels of respiration, confirming the intricate relationships between oro-facial behaviors in mice (Fig. S5).

**Figure 4.**
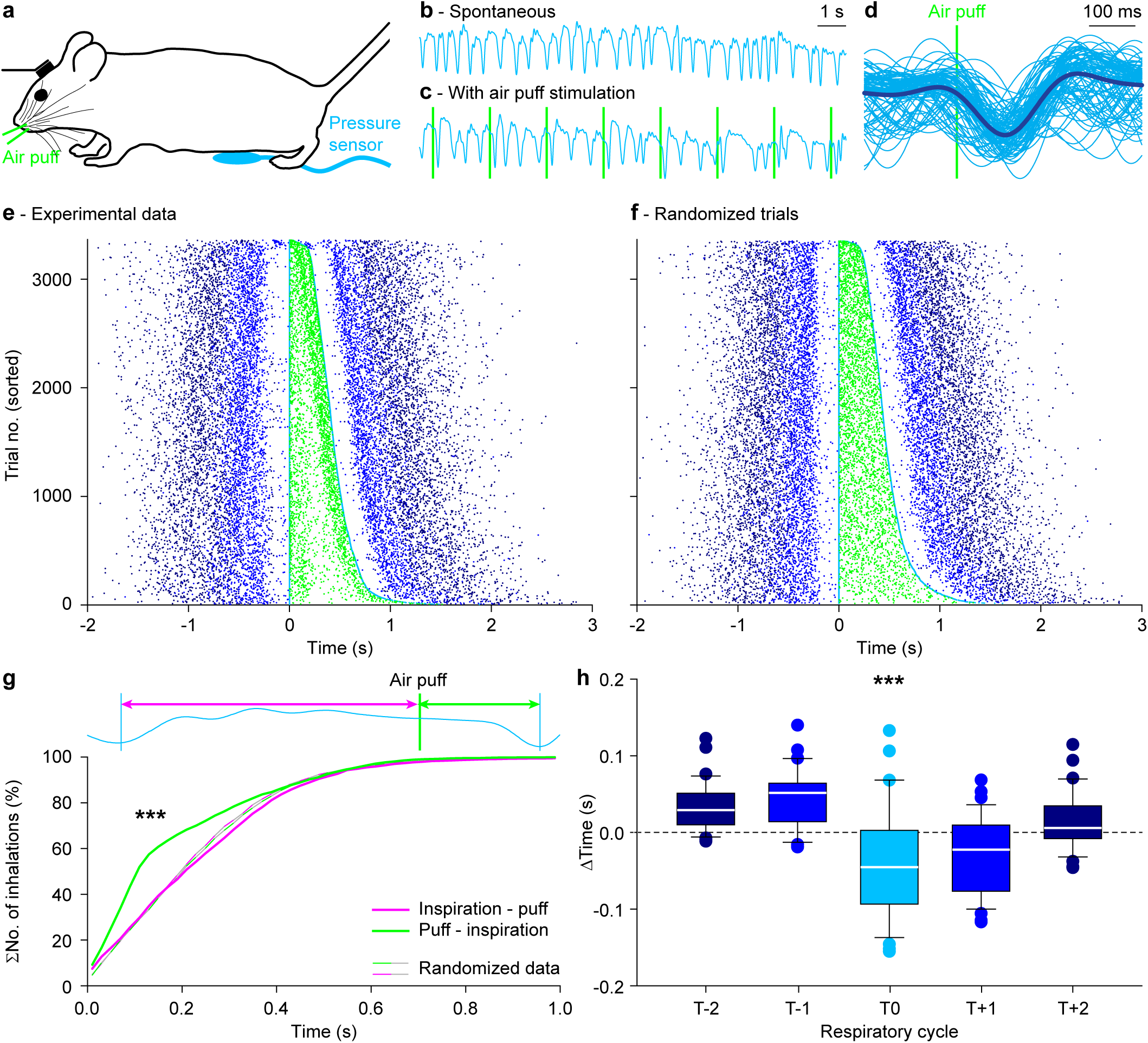
Whisker pad stimulation triggers inspiration. **a** Every 2 s, mice received an air puff to their whisker pad while their respiration was recorded using an abdominal pressure sensor. In comparison to the relatively regular breathing in the absence of air puff stimulation (**b**), the respiratory pattern appeared to be affected by sensory stimulation (**c**). Whisker pad stimulation (vertical lines) often triggered inhalation. **d** The raw respiratory signals around the air puff (90 trials of the same experiment as in **c** with the average (thick line) overlaid) demonstrate that whisker pad stimulation often triggers inspiration. **e** Raster plots showing respiratory cycles from 12 mice pooled together and sorted based upon the duration of the respiratory cycle during which the air puff (light green dots) was applied. The trials were aligned on the start of the last inspiration before the onset of the stimulus. Cyan dots indicate the start of the last inspiration before and the first inspiration after the air puff. The previous and subsequent respiratory cycles are indicated by increasingly darker shades of blue (see color code of **h**). In this plot the air puffs are concentrated just after or just before the start of an inspiration. The latter reflect the triggering of the next inspiration by the air puff. This effect was not observed when the stimulation occurred just after the start of inspiration. **f** Upon random shuffling of the respiratory cycles, the air puffs are equally distributed over the respiratory cycle. **g** Cumulative distributions of inspiration-puff and puff-inspiration intervals for experimental and randomized data. *** *p* < 0.001, Kolmogorov-Smirnov test. **h** Box plots of the duration of the respiratory cycles around the puff indicated that indeed the cycle during which the whisker pad stimulation was given were shorter. T0 is the cycle during which the air puff was given.

### Purkinje cells sensitive for whisker stimulation can contribute to changes in respiratory timing

Given that Purkinje cells in the lateral cerebellum respond to whisker stimulation (Bosman et al., 2010; Brown and Raman, 2018; Romano et al., 2018) and modulate their firing rate along the respiratory cycle, we examined whether whisker-sensitive Purkinje cells could mediate the stimulation-induced change in respiratory timing. To this end, we compared the spiking pattern of 57 Purkinje cells before and after whisker stimulation. Of these Purkinje cells, 53 (93%) responded with a statistically significant complex spike response (Fig. S6). First, we examined the firing pattern during the whole period with 0.5 Hz whisker pad stimulation. In contrast to unperturbed respiration (Fig. 1b), about half of the Purkinje cells showed their peak in complex spike activity during the last quarter of the respiratory cycle, corresponding to expiration (Fig. 5a-c,e). The strongest peak in complex spike firing occurred around 40 ms after whisker pad stimulation (Fig. S7a-b), thus approximately 50 ms before the average start of the first inspiration after the stimulus (Fig. 4e). To test the possibility that the complex spikes accelerate the occurrence of the next inspiration, we compared the timing of complex spike firing during individual trials relative to that of the start of the inspiration. However, we found no clear relation between them (Fig. S7a). Accordingly, when we compared trials with and without complex spikes, we could not find any obvious difference in the timing of the next or subsequent start of inspiration (Fig. S7b-e). Thus, we conclude that the complex spikes observed in the lateral cerebellum, although reacting to whisker stimulation, do not modulate the timing of respiratory responses to whisker pad stimulation in the short-term.

**Figure 5.**
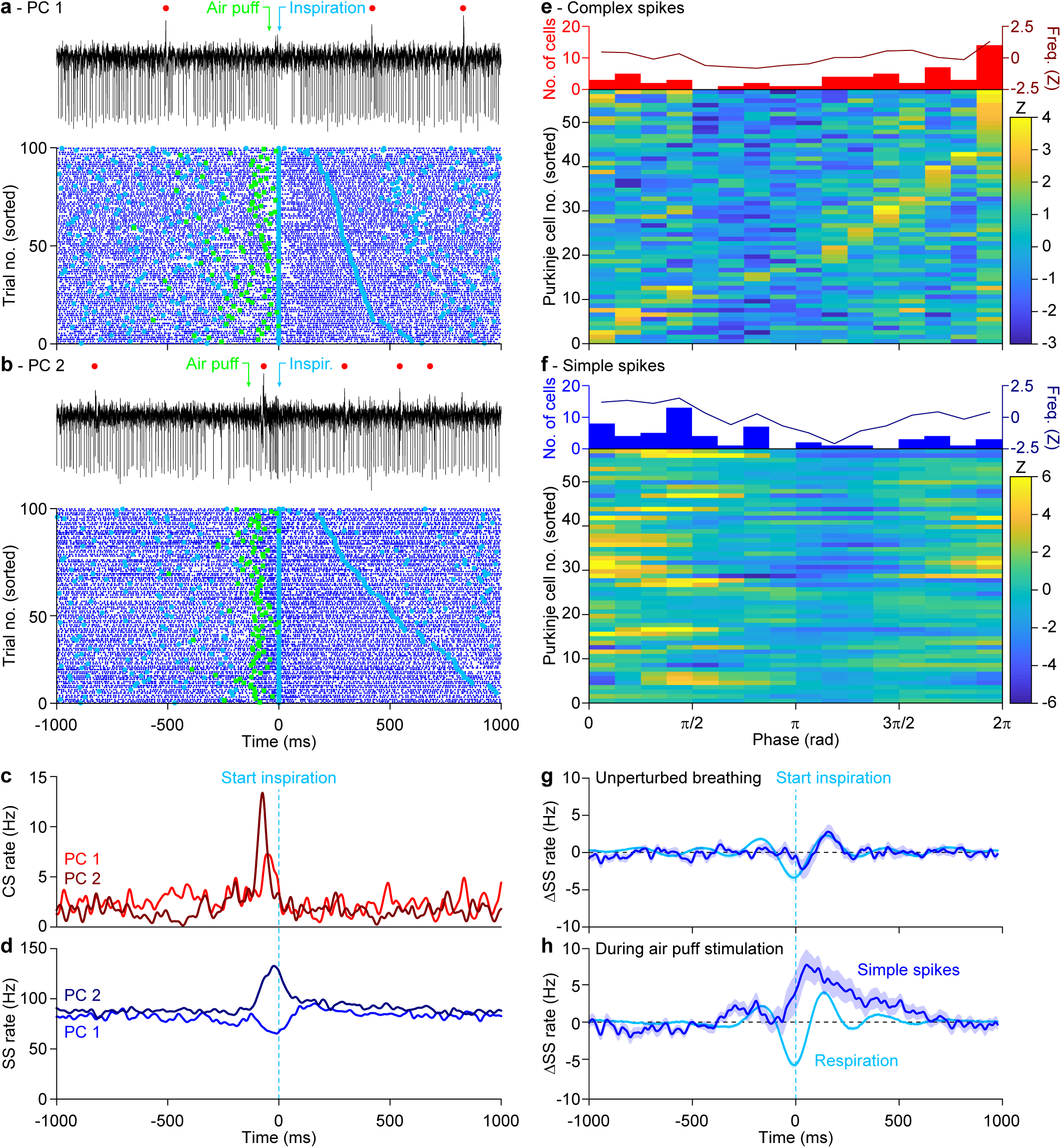
Purkinje cell activity anticipates respiratory responses. Representative Purkinje cell recordings showing either suppressed (**a**) or increased (**b**) simple spike firing upon whisker pad air puff stimulation. Above the trace, the complex spikes are indicated by red dots, the start of the air puff by a green arrow and the start of inspiration by a cyan arrow. Below the traces are raster plots of simple spike activity aligned on the start of the first inspiration after the air puff. In the raster plots, the air puffs are indicated by green squares. The trials are sorted based on the duration of the respiratory interval following the air puff. The starting moment of inspiration are indicated by cyan dots. Note that the complex spikes are not shown in the raster plots for reasons of clarity. Convolved histograms of the complex spikes (**c**) and simple spikes (**d**) of these two illustrated Purkinje cells aligned on the first inspiration onsets after stimulus. The complex spike (**e**) and simple spike (**f**) data of the entire population of 57 Purkinje cells measured in this way are indicated in heat maps. The Purkinje cells are sorted according to the moments of their maximal complex spike firing. In **g** is illustrated the same plot of Fig. 2e for comparison along with the homologous plots for the air puff induced anticipated inhalations (**h**). Even in this case it can be observed a similarity between the profiles of the averaged respiratory signal and the averaged simple spike activity. In the latter case the simple spikes modulation anticipate the averaged respiration signal.

As detailed in a previous publication (Romano et al., 2018), simple spike firing was also considerably affected by whisker pad stimulation (Fig. 5a-b,d; Fig. S6). Likewise, the temporal relation between simple spike and complex spike firing, as found during unperturbed respiration (Fig. 1b,d; S2b-c), was disrupted and no longer significant during the whole period with 0.5 Hz whisker stimulation (p > 0.05 for simple spike firing in all bins compared to the bin with the strongest complex spike modulation; Pearson correlation tests with Benjamini-Hochberg correction for multiple comparisons) (Fig. 5e-f). When we related simple spike modulation to the respiratory rhythm following whisker stimulation, we found that 20 out of the 32 (62%) Purkinje cells with a significant simple spike modulation predominantly increased their simple spike activity, whereas 12 (38%) predominantly decreased their simple spike firing. Compared to unperturbed respiration (Fig. 5g), the population increase of simple spike firing now peaked during earlier phases of the respiratory cycle, pointing towards an acceleration in their activity (*p* = 0.034, Wilcoxon test; Fig. 5h). Restricting the analysis to the cycle around the air puff, it became apparent that the population average of simple spike firing now preceded the change in respiratory behavior, suggesting that the air puff-triggered simple spike response could contribute to the observed acceleration of inspiration. We further examined this possibility by performing a trial-by-trial analysis of variation. During eupneic breathing the prevalence of correlation was below the 45° line for both suppressive and facilitating Purkinje cells, indicating that under these circumstances the simple spike modulation follows respiration and therefore cannot control it (Fig. 3). However, during perturbed respiration, the modulation of simple spike firing preceded the ongoing respiration by a few tens of milliseconds (Fig. 6). Moreover, when we segregated the cells that showed simple spike modulation to whisker stimulation (Fig. 6c) from those that did not (Fig. 6d), we observed that the maximal correlations between respiration and simple spike firing were on average stronger in the whisker-related Purkinje cells (r = 0.35 ± 0.08 vs. 0.28 ± 0.05; *p* = 0.018, t = 2.498, df = 31, t test). These data confirm that simple spike responses following whisker stimulation are endowed with the temporal features for accelerating respiratory responses, whereby the simple spike responses predict the strength of the inhalation.

**Figure 6.**
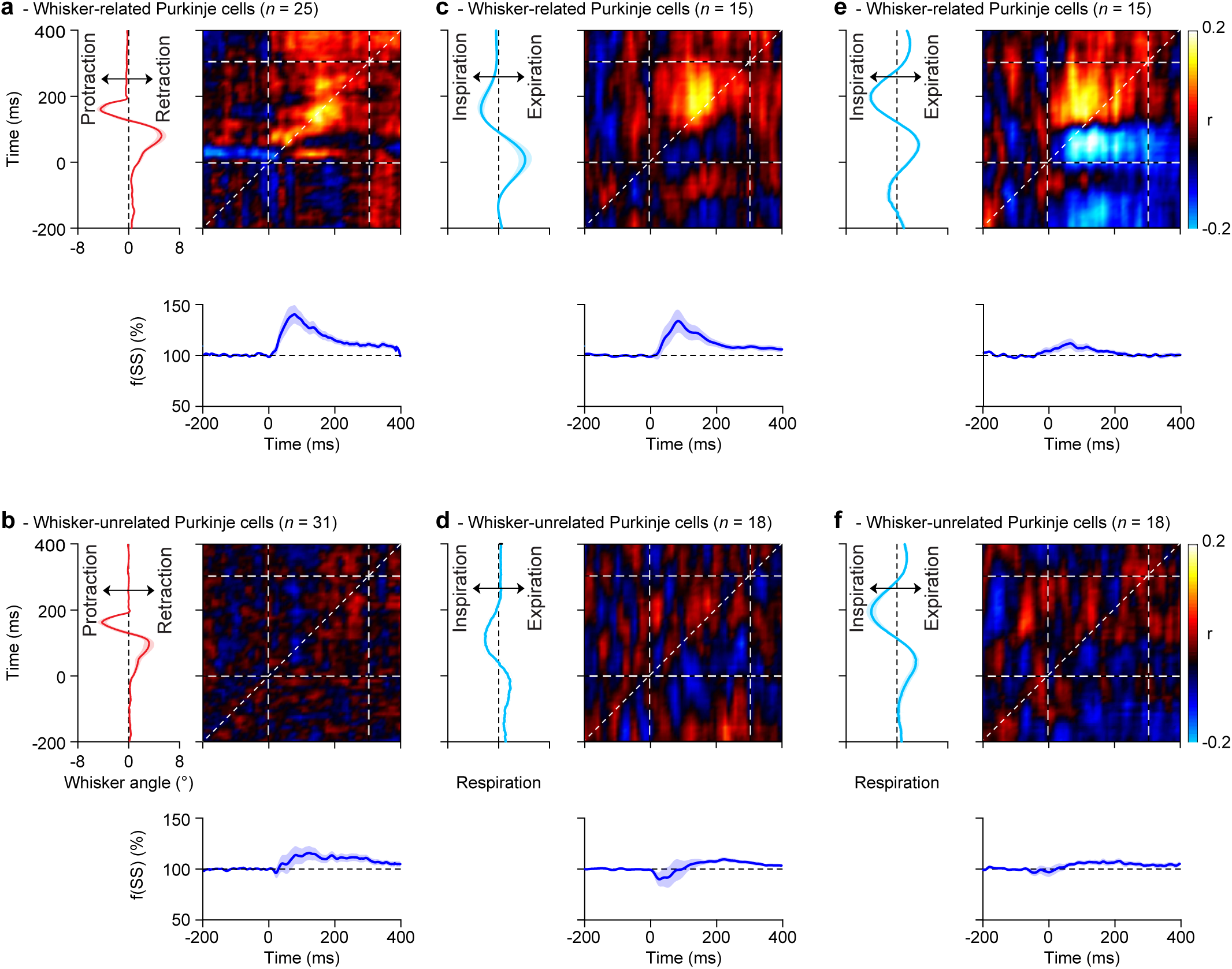
Modulation of simple spike firing precedes respiratory adaptation in whisker-related Purkinje cells. Correlation matrix between simple spike firing (blue trace at the bottom represents the averaged convolved peri-stimulus time histograms) and mean reflexive whisker protraction (red trace on the left) for Purkinje cells with (**a**, *n* = 25) and without (**b**, *n* = 31) significant correlation between simple spike firing and air puff-induced whisker movement (see Methods). For 15 out of the 25 whisker-related cells and 18 out of 31 whisker-unrelated cells the respiratory signal was simultaneously recorded and used for the respiration-spike matrix of correlation in **c** and **d**, respectively. The whisker-related Purkinje cells had a higher correlation between their instantaneous simple spike rate and respiration than the other Purkinje cells (*p* = 0.018, *t* test). The location of the maximal correlation above the 45° line indicates that in trials in which the Purkinje cells fired more simple spike then, few tens of milliseconds later, the amplitude of the respiration was bigger and vice versa. In addition, the simple spike to whisker correlation (for the whisker-related cells) is stronger and earlier in time when the matrix of correlation is aligned to the puff-induced inhalation, rather than to the puff itself (**e**). Conversely, on average the whisker-unrelated cells did not show a clear correlation even when the signals were aligned to the air puff-induced inspiration (**f**). Shaded areas around the traces indicate SEM.

### Respiration related Purkinje cells are located in specific portions of the cerebellar cortex

Next, we mapped the location of the Purkinje cells recorded in this study. During unperturbed respiration, the strongest complex spike modulation was found laterally in crus 2 (Fig. 7a; left column). This complex spike hot spot extended rostrally into crus 1 during respiration perturbed by whisker stimulation (Fig. 7b). The Purkinje cells in crus 1 that were recruited during perturbed, but not during unperturbed respiration, were mainly those that responded with a complex spike response directly to the sensory stimulation (Fig. 7c). The simple spike responses showed a distribution that was largely complementary to that of the complex spike responses (Fig. 7; right column). The Purkinje cells with a predominantly increased simple spike rate during an unperturbed respiratory cycle were largely found around the border between the vermis and the simple lobule and crus 1, with a few cells extremely lateral in crus 1 (Fig. 7a). The cells that showed decreased simple spike firing during unperturbed respiration were largely confined to a parasagittally oriented strip in the middle of crus 1 and crus 2. During perturbed respiration, this pattern was largely unaltered, although the lateral regions now also showed a decreased firing rate (Fig. 7b). Importantly, the Purkinje cells in medial crus 1 showed particularly strong correlations to both whisker inputs and respiration.

**Figure 7.**
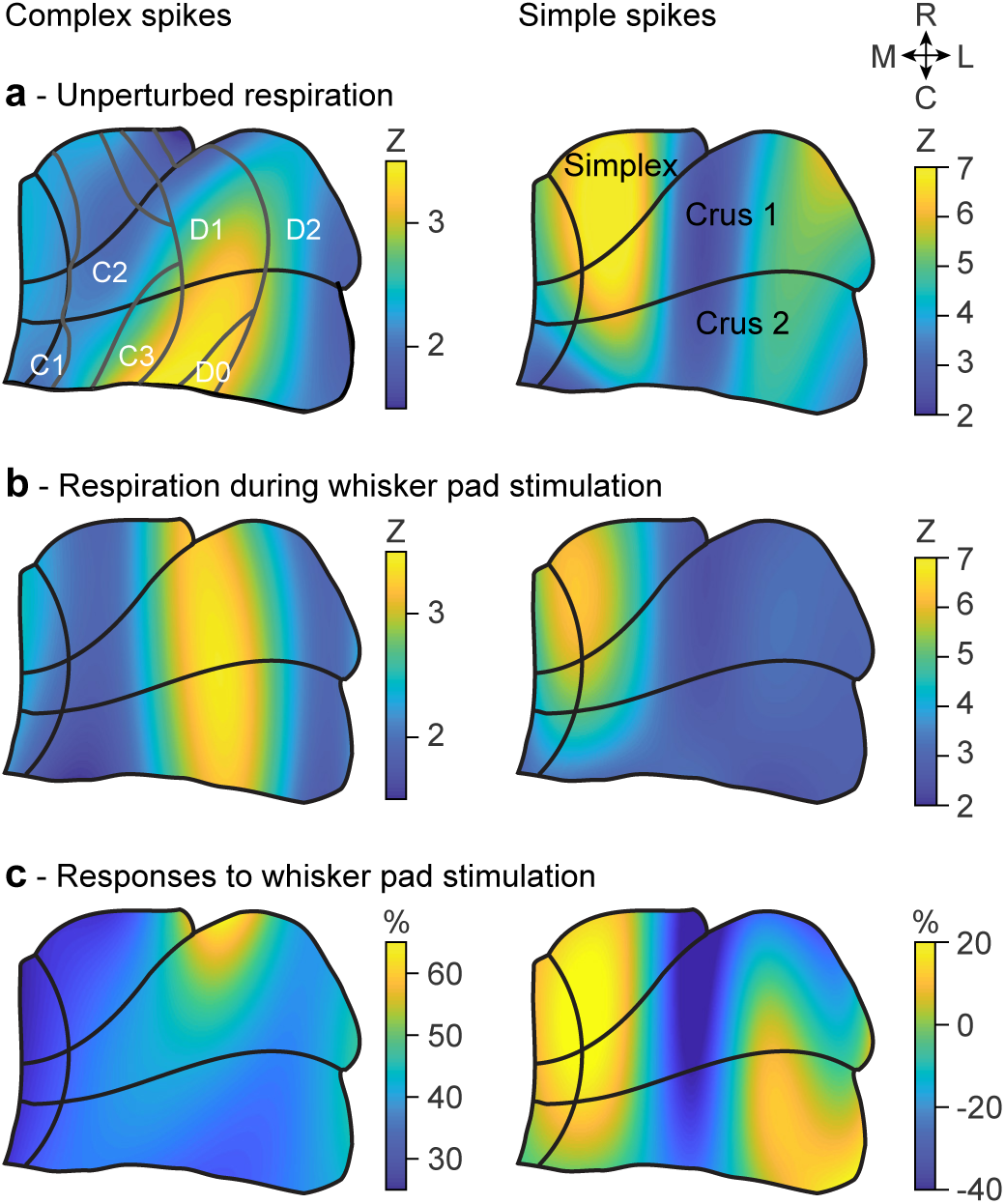
Respiration related Purkinje cells are located in specific portions of the cerebellar cortex. **a** Based upon the entry point of each electrode relative to a standardized map of the simplex, crus 1 and crus 2 lobules, a map was created indicating the spatial distribution of the maximal increase (in Z score) of complex spikes (left) and simple spikes (right) during the respiratory cycle in the absence of whisker pad stimulation. **b** The same during the presence of whisker pad stimulation. This analysis revealed an area with a relatively strong correlation between respiration and complex spike firing in the medio-lateral part of crus 2 extending rostrally in crus 1. Simple spikes correlated to respiration were mainly found medially in the simple lobule and crus 1. There were some differences in the spatial pattern of responses during unperturbed and perturbed respiration. These could be partially explained by the pattern of response probabilities (in percentage of baseline firing) to whisker pad air puff stimulation (**c**). In the left panel of **a**, the tentative locations of the cerebellar modules are indicated. C = caudal, L = lateral, M = medial, R = rostral.

### Purkinje cell stimulation mimics the impact of whisker stimulation on respiratory timing

As our analyses revealed that during perturbed respiration increased simple spike firing preceded the accelerated inspiration, we wondered whether we could mimic the impact of whisker pad air puff stimulation on the timing of inspiration by transiently stimulating the Purkinje cells in the medial parts of lobule simplex as well as the crus 1 and crus 2 areas highlighted above. To this end, we made use of transgenic mice that expressed channelrhodopsin specifically in their Purkinje cells (Pcp2-Ai27 mice) (Witter et al., 2013; Romano et al., 2018). In line with previous whole cell recordings in vivo (Witter et al., 2013), a brief pulse of blue light triggered a strong increase in simple spike firing (Fig. 8a-b). Purkinje cell optogenetic stimulation significantly accelerated the occurrence of the next respiratory cycle (Fig. 8c-d). The inspiration started 189 ms (median value, IQR: 243 ms) after the onset of optogenetic stimulation compared to 224 ms (IQR: 212 ms) during the sham-control condition. Consequently, inspiration was accelerated due to optogenetic Purkinje cell stimulation (*p* < 0.001, Kolmogorov-Smirnov test), but not in the sham-control experiments (*p* = 0.220, Kolmogorov-Smirnov test; Fig. 8d). We conclude that optogenetic stimulation of the Purkinje cell activity in lobule simplex as well as crus 1 and crus 2 areas is sufficient to induce an acceleration in the occurrence of the next respiratory cycle.

**Figure 8.**
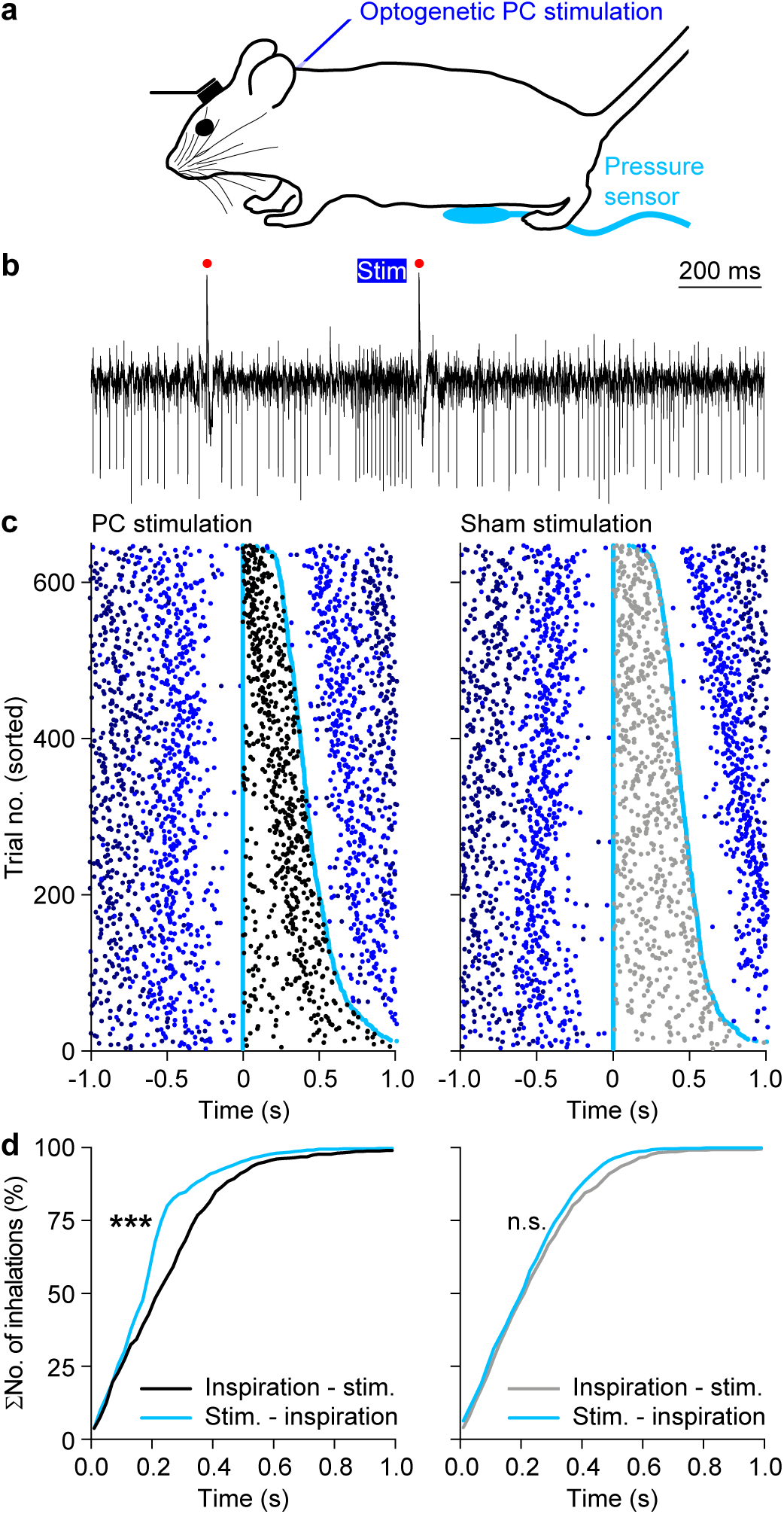
Optogenetic stimulation of Purkinje cells triggers accelerated inspiration. **a** The impact of optogenetic stimulation on respiratory timing was studied using transgenic mice expressing ChR2 exclusively in their Purkinje cells (PCs). **b** 100 ms light pulses caused brief increases in simple spike firing. Red dots indicate complex spikes. **c** Raster plots showing respiratory cycles from 13 mice pooled together and sorted based upon the duration of the respiratory cycle during which optogenetic Purkinje cell stimulus (black dots, left) or the sham stimulus (grey dots, right) was applied. The trials were aligned on the start of the last inhalation before the onset of the stimulus. Cyan dots indicate the start of the last inspiration before and the first inspiration after the stimulus. The previous and subsequent respiratory cycles are indicated by increasingly darker shades of blue. Inspiration typically started around 200 ms after the onset of Purkinje cell stimulation. **d** Cumulative histograms of the interval between the start of inspiration and the stimulus (grey / black) and between the start of the stimulus and the start of the next inspiration (cyan). Purkinje cell stimulation (left), but not sham stimulation (right), accelerates the start of the next inspiration. *** *p* < 0.001, Kolmogorov-Smirnov test.

### Modification of AMPA receptors at parallel fiber to Purkinje cell synapse impairs impact of whisker stimulation on respiratory timing

To find out whether functionally intact cerebellar Purkinje cells are necessary for the respiratory changes induced with whisker stimulation, we investigated this response in a mouse model that lacked the AMPA GluA3 subunit at their parallel fiber to Purkinje cell synapses (Gutierrez-Castellanos et al., 2017). This mutant (Pcp2-Gria3^−/−^ mice) has been shown to be impaired in its simple spike modulation following whisker stimulation (Romano et al., 2018) (Fig. 9a). The Pcp2-Gria3^−/−^ mice showed a normal frequency of respiration during unperturbed respiration (2.6 ± 0.5 Hz versus 2.7 ± 0.8 Hz for wild type mice; *p* = 0.713, t = 0.0.378, df = 10, t test) (Fig. 9b). However, the mutants were significantly impaired in their ability to advance the respiratory response following whisker stimulation (*p* < 0.001, Kolmogorov-Smirnov test) (Fig. 9c-e). Moreover, the simple spike firing of their Purkinje cells was significantly different from that in the wild types in that they did not precede respiration directly following whisker stimulation (*p* = 0.040, Pcp2-Gria3^−/−^ versus wild type, Wilcoxon Rank Sum test) (Fig. 9f-g). These data indicate that a cerebellar, cell-specific interference with a mechanism that has the potential to increase the simple spike firing rate results in a hampered ability to accelerate the respiratory response.

**Figure 9.**
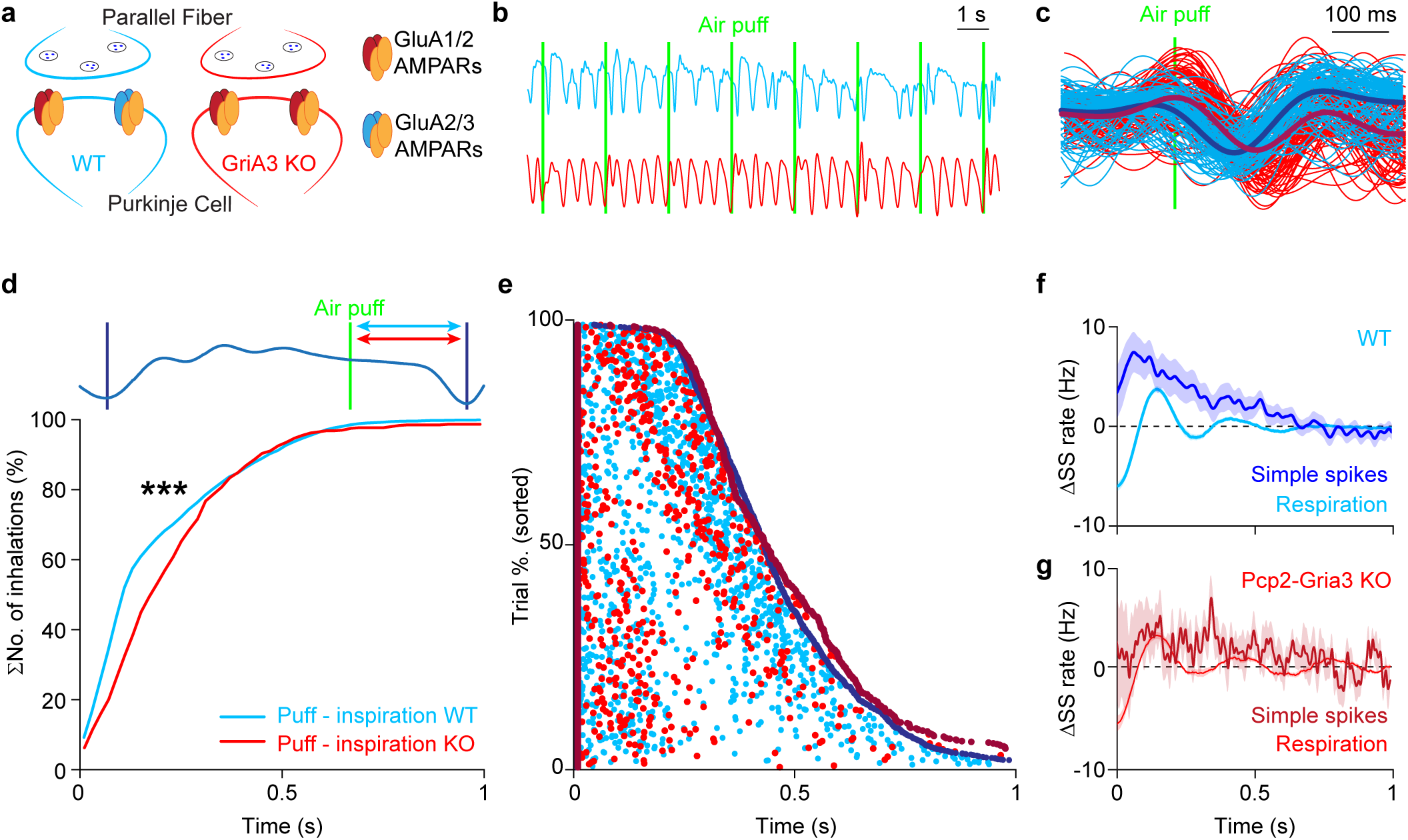
Delayed respiratory response in Gria3 KO mice. **a** Pcp2-Gria3 KO mice lack the gene for AMPA receptor GluA3 subunit, specifically in Purkinje cells. Instead, they express more GluA1/GluA2-type AMPA receptors than wild type mice. **b** In Pcp2-Gria3 KO mice, the whisker stimulation (vertical green lines) appears to be less effective in triggering inhalation than in wild type mice. **c** The respiratory pattern of an exemplary Gria3 KO mouse (red lines) around the air puff showing delayed inspiration when compared to a wild type mouse (the cyan lines are the same as in Fig. 4c-d). The puff-inspiration intervals of 6 Gria3 KO mice are longer compared to those of wild type mice (*p* < 0.001, Kolmogorov-Smirnov test). This results in a shift to the right of their cumulative histogram (**d**). In **e** the raster plot of the respiratory cycles perturbed by the air puffs are sorted from the shortest (top) to the longest (bottom). The beginning and the end of each cycle are represented with dark blue dots for the wild type and dark red dots for the Pcp2-Gria3 KO mice, while the relative time of the air puffs is depicted in cyan and red respectively. Looking at all the individual data points, both red and cyan dots are not randomly distributed and tend to accumulate before the subsequent inhalation. The delay to the start of the next inspiration is longer in Pcp2-Gria3 KO mice than in wild type mice. The simple spike activity of intact Purkinje cells increases during the air puff-triggered inspiration (**f**). This increase resamples, precedes and potentially affect the relative ongoing respiration. Conversely, the simple spike activity of the Pcp2-GriA3 KO mice modulates differently than in WT mice (*p* < 0.001, Kolmogorov-Smirnov test) and do not resample the respiration signal (**g**).

## Discussion

Animals use periodic and oscillatory behaviors in a variety of functional movements and they display adaptation in the coordination of such behaviors as a function of systematic changes of the environment. For instance, during sniffing rodents coordinate the movements of their whiskers with those of the respiratory and olfactory system (Welker, 1964; Kurnikova et al., 2017) and this behavior can be adjusted to the discrimination task involved. The way the brain organizes such control mechanisms is largely unknown. Here we investigated the involvement of the cerebellum in the whisker and respiratory systems by recording, stimulating and impairing Purkinje cells in the lateral cerebellum where respiration and whisking converge. During eupneic respiration, Purkinje cells in the simplex, crus 1 and crus 2 lobules of the cerebellum fire in tune with specific phases of the respiratory rhythm. During such unperturbed breathing, we found no evidence for a motor control function of either the complex spikes or the simple spikes. The response could be described rather as an efference copy of the respiratory signal, modulated by specific phases of the respiratory rhythm. In contrast, stimulation of the whiskers with air puffs accelerated simple spike activity that in turn contributed to accelerated periods of inspiration. These increases in simple spike activity in the lateral cerebellum are probably sufficient to drive and integrate different forms of rhythmic behavior, because increases in simple spike activity that were artificially evoked by optogenetic stimulation of Purkinje cells also accelerated the respiratory cycle in a phase-dependent manner. Finally, cell-specific blockage of AMPA receptor signaling in Purkinje cells impaired the ability of the mice to advance both their simple spike response and their respiratory response following air puff stimulation of the whiskers, highlighting the necessity of an intact cerebellum for synergistic phase control.

Rhythmic behaviors are part of everyday life of mammals, they are modified depending on ongoing behavioral demands, and they are often synergistically adjusted to warrant optimal coordination (Moore et al., 2013; Kurnikova et al., 2017). It is known that different rhythmic behaviors can be controlled as muscle synergies by local networks. For instance, coordination between the left and right hind-limb can still take place during locomotion even after the descending connections from the motor cortex and brainstem to the spinal cord are disrupted (Desrochers et al., 2019). Likewise, when cerebellar function is impaired, the basic muscle activities of breathing and related orofacial behaviors can still take place up to a degree (Chen et al., 2005; Abecassis et al., 2017; Bellavance et al., 2017). Thus, local networks in the spinal cord and the brainstem are sufficient to generate the basic antagonistic muscle activities that mediate the rhythmic properties of locomotion and breathing, respectively (Tresch et al., 1999; Talpalar et al., 2013; Bellavance et al., 2017; Kurnikova et al., 2017). However, the cerebellum may well play an instrumental role in coordinating front and hind limb movements during more elaborate forms of locomotion adaptation (Hoogland et al., 2015; Machado et al., 2015; Vinueza Veloz et al., 2015; Darmohray et al., 2019) or adjusting the respiratory cycle during more complex tasks such as speech (Deger et al., 1999). Here, we provide evidence that changes in simple spike activity of Purkinje cells in the simplex, crus 1 and crus 2 areas, in which respiratory and whisking processing converges, may contribute to re-adaptation of the respiratory timing signal following sensory perturbation of the facial whiskers. Natural or artificial activation of Purkinje cells sensitive for whisker stimulation can accelerate the occurrence of the next inspiration. Interestingly, increases in simple spike activity of the same Purkinje cells can also drive and accelerate whisker protraction (Fig. 6), allowing for synergistic control of two different forms of orofacial behaviors.

To the best of our knowledge, this is one of the first studies indicating that the simple spike activity of individual Purkinje cells can simultaneously drive different forms of motor behavior, in this case the rhythmic behaviors represented by breathing and whisking. This finding elaborates on several behavioral studies demonstrating the role of the cerebellum in synergistic control of diverse motor domains. For example, the olivocerebellar system has been shown to be involved in the coordination between eye and hand movements (Kitazawa et al., 1998; Vinueza Veloz et al., 2015; Owens et al., 2018), between trunk and limb movements (Bakker et al., 2006; Caliandro et al., 2017), as well as between shoulder, arm and finger movements (Thach et al., 1993; Timmann et al., 2000). The current study amasses to that lot by demonstrating for the first time functional convergence of autonomic and sensorimotor behaviors on single Purkinje cells. We demonstrate that the advancement of the respiratory cycle is related to anticipatory Purkinje cell responses in the same cells that signal the reflexive whisking behavior. Given their rich and diverse parallel fiber inputs mediating signals from different sensorimotor systems, and given the well-timed and powerful climbing fiber input that can trigger plasticity (Ito, 2000; Coesmans et al., 2004; De Zeeuw et al., 2011; Gao et al., 2012), we postulate that Purkinje cells and the cerebellar cortex as a whole mediate synergy and integration of different motor domains in both voluntary and autonomic systems.

The contribution of complex spikes to the acute changes in respiratory behavior remains to be elucidated. The prominent complex spike response at a particular phase of the respiratory rhythm induces plasticity at particular phases of respiration, suggesting a primary role in binding different synergies. Nevertheless, acute respiratory responses during trials with and without complex spikes did not reveal any significant difference in behavior. This does not exclude the possibility that another system is being modulated by the complex spike signal. We also could not find any differential role for complex spikes during trials with and without air puff stimulation. This means that the phase related signal conveyed by the complex spike is robust to the sensory perturbation. The only prominent difference that occurred in the trials with unperturbed and perturbed breathing is that the interval between peak activity of the simple spikes and that of the complex spikes robustly changed in the trials with air puff stimulation. This difference may well have impact on the induction of various forms of cellular plasticity in the cerebellar cortex that depend on the moment of climbing fiber activation with respect to the baseline activity in the Purkinje cells and network involved (Wang et al., 2000; Coesmans et al., 2004; Gao et al., 2012; De Zeeuw and Ten Brinke, 2015; Suvrathan et al., 2016). Indeed, this possibility agrees with the fact that the complex spike frequency negatively correlates with the induction of long-term potentiation (LTP) (Coesmans et al., 2004). It is also consistent with the observation that ablating this form of plasticity in the Pcp2-Gria3^−/−^ mice corrupted the synergistic behavioral response following whisker stimulation. It will be interesting to investigate to what extent an induced shift in complex spike phase would have an impact on the relationship between different rhythms, including that of respiration.

It is likely that Purkinje cells in the cerebellar cortex influence the nuclei in the brainstem that control breathing and/or whisking. The cerebellar fastigial nucleus is known to modulate the respiratory cycle by sensing O_2_ and CO_2_ in the blood (Xu and Frazier, 2000; Xu et al., 2001; Martino et al., 2006; Martino et al., 2007). The roles of the other cerebellar nuclei, which do not seem to have correlates of pH sensors (Xu et al., 2001), are still controversial (Xu and Frazier, 2000). Possibly, the interposed nuclei play a role in control of the upper airways, as bilateral lesions of this region suppress coughing responses (Xu et al., 1997). The central pattern generator for respiration is located in the preBötzinger complex (Smith et al., 1991; Ramirez et al., 2011; Feldman et al., 2013; Moore et al., 2013). There is no direct connection from the cerebellar nuclei to the preBötzinger complex nor to the adjacent Bötzinger complex (Teune et al., 2000), which is consistent with our finding that the Purkinje cell-mediated impact of whisker stimulation changes the timing, not the frequency of respiration (Fig. 4h). Although the anatomical pathways via which the cerebellum could affect respiration are still matter of debate, projections to nuclei downstream of the preBötzinger complex are expected. Possibly, the projection to the region of the post-inspiratory complex (Anderson et al., 2016) at the border of the intermediate and gigantocellular reticular formation close to the ambiguus nucleus (Teune et al., 2000; Lu et al., 2013) plays a role, which would be in line with the increased Purkinje cell activity found during post-inspiration. Also the direct connection to the parabrachial complex (Teune et al., 2000), which in turn projects to the phrenic nucleus in the spinal cord that houses the motor neurons of the diaphragm, cannot be excluded (Dobbins and Feldman, 1994). The projections of the cerebellar interposed and lateral nuclei, that mediate the Purkinje cell output of the lateral cerebellar cortex, to the brainstem nuclei downstream of the central pattern generator suggest that the lateral cerebellum preferentially affects the timing, not the frequency, of respiration, which is in line with our experimental data.

We conclude that individual Purkinje cells in the lateral cerebellum can synergistically coordinate multiple motor behaviours, such as respiration and whisking, by inserting accelerations into the cerebellar sensorimotor studies.

## Methods

### Mice

We used 18 WT mice with a C57BL/6J background (13 males and 5 females from Charles Rivers, Leiden, the Netherlands) for the electrophysiological recordings and compared their behavior to 13 Tg(Pcp2-cre)2Mpin;Gt(ROSA)26Sor^tm27.1(CAG-OP4*H134R/tdTomato)Hze^ KO mice (Witter et al., 2013) expressing channelrhodopsin-2 for optogenetic stimulation of their Purkinje cells (6 males and 7 females from the same breeding colony as the WT mice, preferably using littermates). In addition, we used 6 Tg(Pcp2-cre)2Mpin;Gria3^tm2Rsp^ KO mice (Gutierrez-Castellanos et al., 2017)nuclei, also on a C57BL/6J background. The mice had an age of 4-7 months. Mice were socially housed until surgery and single-housed afterwards. The mice were kept at a 12/12 h light/dark cycle and had not been used for any other study before the start of the experiments described here. All experimental procedures were approved a priori by an independent animal ethical committee (DEC-Consult, Soest, The Netherlands) as required by Dutch law and conform the relevant institutional regulations of the Erasmus MC and Dutch legislation on animal experimentation. Permission was filed under the license numbers EMC3001 and AVD101002015273.

### Surgeries

All mice received a magnetic pedestal that was attached to the skull above bregma using Optibond adhesive (Kerr Corporation, Orange, CA) and a craniotomy was made on top of crus 1 and crus 2. The surgical procedures were performed under isoflurane anesthesia (2-4% V/V in O_2_). Post-surgical pain was treated with 5 mg/kg carprofen (“Rimadyl”, Pfizer, New York, NY), 1 µg lidocaine (Braun, Meisingen, Germany), 50 µg/kg buprenorphine (“Temgesic”, Reckitt Benckiser Pharmaceuticals, Slough, United Kingdom) and 1 µg bupivacaine (Actavis, Parsippany-Troy Hills, NJ, USA). After three days of recovery, mice were habituated to the recording setup during at least 2 daily sessions of approximately 45 min. In the recording setup they were head-fixed using the magnetic pedestal. Further body movements were prevented by using a customized restrainer and filling the empty space with paper tissues.

### Respiration recordings and phase transformation

Respiration was recorded using a PowerLab 4/30 analog-to-digital converter (AD Instruments, Oxford, United Kingdom) in combination with a pressure sensor that was placed at the abdomen of the mice. The resulting signal was filtered with MATLAB’s (MathWorks, Natick, MA, USA) Butterworth bandpass filter (cutoff frequencies 1 and 10 Hz, chosen to include respiratory frequencies visible on the Fourier transform of the raw signal). For the averages of the respiration signal around the stimulus, movement artefacts were removed by excluding trials in which the signal surpassed three times the standard deviation in a 200 ms window before the stimulus. The phase transform of the respiration signal was acquired with the co_hilbproto (which calculates a ‘protophase’ of a scalar time series using the Hilbert transform) and co_fbtrT (protophase to phase transformation) functions from MATLAB toolbox DAMOCO. As the default setting, the DAMOCO toolbox chooses as initial phase the maximum of the respiration signal, but for our analysis it was more beneficial to set zero phase at the moment when the mouse starts inspiration. Therefore, before the phase transform the respiration signal was multiplied by −1, so that no changes needed to be made to the functions of this toolbox.

### Whisker pad stimulation and whisker tracking

Sensory stimulation (0.5 Hz) was given to the center of the whisker pad of awake mice by means of air puffs given from approximately 5 mm at 30 degree angle with the whisker pad. Each puff was around 2 bar and had a duration of 30 ms. Videos of the movements of the untrimmed large facial whiskers were made from above using a bright LED panel as back-light (λ = 640 nm) at a frame rate of 1,000 Hz (480 x 500 pixels using an A504k camera from Basler Vision Technologies, Ahrensburg, Germany). The whisker movements were tracked as described previously (Rahmati et al., 2014; Romano et al., 2018) using the BIOTACT Whisker Tracking Tool (Perkon et al., 2011) in combination with custom written code (https://github.com/elifesciences-publications/BWTT_PP). The whisker movements were described as the average angle of all trackable whiskers per frame.

### Electrophysiology

Electrophysiological recordings were performed in awake mice using quartz-coated platinum/tungsten electrodes (2-5 MΩ, outer diameter = 80 µm, Thomas Recording, Giessen, Germany). The latter electrodes were placed in an 8×4 matrix (Thomas Recording), with an inter-electrode distance of 305 µm. Prior to the recordings, the mice were lightly anesthetized with isoflurane to remove the dura, bring them in the setup and adjust all manipulators. Recordings started at least 60 min after termination of anesthesia and were made in lobules simplex, crus 1 and crus 2 ipsilateral to the side of the whisker pad stimulation at a minimal depth of 500 µm. The electrophysiological signal was digitized at 25 kHz, using a 1-6,000 Hz band-pass filter, 22x pre-amplified and stored using a RZ2 multi-channel workstation (Tucker-Davis Technologies, Alachua, FL). Spikes were detected offline using SpikeTrain (Neurasmus, Rotterdam, The Netherlands). A recording was considered to originate from a single Purkinje cell when it contained both complex spikes (identified by the presence of stereotypic spikelets) and simple spikes, when the minimal inter-spike interval of simple spikes was 3 ms and when each complex spike was followed by a pause in simple spike firing of at least 8 ms. Only single-unit recordings of Purkinje cells with a minimum recording duration of 120 s were selected for further analysis.

### Optogenetic stimulation

LED photostimulation (λ = 470 nm) driven by a Thorlabs LED driver (225 µW) was given through an optic fiber (400 µm in diameter, Thorlabs, Newton, NJ, USA). The optic fiber rested on the dura mater above the midline between crus 1, crus 2, approximately 1 mm lateral from the vermis, via the craniotomy.

### Polar plots

Polar plots were generated to describe the correlation between respiratory phase and Purkinje cell spiking activity. To this end, we attributed each spike to a phase (using 16 bins) of the respiration and we compared the recorded distribution with a bootstrap analysis based upon a re-sampling of the spike times after shuffling the inter-spike intervals. The bootstrap analysis was repeated 500 times after which the 99% confidence interval was established. The Z score of each bin was derived by dividing, for each bin, the dividing the difference between number of spikes of a Purkinje cell during that bin and the average number of spikes of all bins by the standard deviation of all bins.

### Trial-by-trial correlation analysis

The inter-trial variations between the respiratory signal and the instantaneous simple spike firing rate (Figs. 3 and 6) or the average whisker angle (Figs. 6, S5) were calculated and represented according to a previously published method (Ten Brinke et al., 2015; Romano et al., 2018). Briefly, during each trial, the filtered respiration signal (see above) was compared to either the instantaneous simple spike rate or the relative whisker position without alignment to the baseline. The instantaneous simple spike rate was obtained by convolving spike occurrences across 1 ms bins with an 8 ms Gaussian kernel. The inter-trial variations were subsequently described by creating a matrix of Pearson correlation values for each 10 x 10 ms bin and visualized as heat maps. In Fig. 6, we separated between those Purkinje cells that had a significant correlation between their instantaneous simple spike rate and the whisker angle and those that had not. Significance was established by testing whether the correlation along the 45° line exceed the 99% confidence interval of a bootstrapped dataset in which the inter-spike times were randomly shuffled 500 times.

### Sorted raster plots

To visualize the relation between stimulation and respiration (Figs. 4e, 8c, 9e and S4) or between complex spike firing and respiration (Fig. S7a), sorted raster plots were constructed. For each plot, a dataset composed of a balanced number of trials of all mice was generated and sorted based upon the duration of the respiratory cycle during which the stimulus was applied (Figs. 4e, 8c, 9e) or that of the respiratory cycle preceding the stimulus presentation (Fig. S4b). The trials presented in Fig. S7a were sorted based upon the interval from the air puff stimulus to the first complex spike following that moment. In Fig. S7a, some of the Purkinje cells were recorded simultaneously, leading to a larger number of trials than in the other plots that are based upon mice. The experimental data were compared to a random shuffling of the durations of the respiratory cycles within each experiment (Fig. 4f). During optogenetic stimulation, trials with and without stimulation were randomly intermingled and the sham stimuli served as control (Fig. 8c). The differences between experimental and control data were substantiated by comparing the distribution of the intervals between the start of the last inspiration prior to stimulation and the moment of stimulation with the distribution of intervals between the stimulation and the start of the subsequent inspiration.

### Anatomical maps

To visualize the distribution of the spike-respiration correlation respiration throughout the lobules simplex, crus 1 and crus 2 we developed an anatomical map of the distribution of the Z-score values obtained by the polar plots (Fig. 7a-b). Since the electrophysiological recordings were performed using a grid of 8 x 4 electrodes (placed always on the same type of craniotomy), we could retrieve the approximate location of each cell and plot the corresponding maximum Z-score on a 8 x 4 matrix. Linear interpolation was used to smooth the edges of adjacent patches and the Matlab function “imagesc” was eventually used to obtain the heating map that was overlapped to a schematic draw of the craniotomy. Similarly, also the air puff responses could be represented by plotting the values of maximum variation of firing rate of each cell (Fig. 7c).

### Data inclusion criteria

Only electrophysiological recordings for which the amplitude and the width of the spikes were constant over time were considered for correlation with the respiratory signal. The recordings in which the amplitude or the width of more three consecutive simple spikes exceeded three standard deviations above or below their average were considered unstable and excluded. In this way, any change in spike rate due to the instability of the recordings was avoided.

### Statistics and visualization

Data were tested for normal distribution using a one-sample Kolmogorov-Smirnov or Shapiro-Wilk test. Parametric tests were used for normally distributed data and non-parametric tests for other data. Whenever applicable, two-sided tests were used. Unless stated otherwise, data are summarized as average ± s.d..

Box plots (e.g., see Fig. 4h) indicate the distribution of the data with the box indicating the interquartile-range around the median (horizontal line). The whiskers indicate the 10^th^ and 90^th^ percentiles. Data points outside the 10^th^-90^th^ percentile range are indicated as separate dots. Violin plots (e.g., see Fig. S1) indicate the distribution of all data points as dots. The contours indicate a convolved histogram of the data points (along the y axis, using a Gaussian kernel and reflected along the vertical axis) and the horizontal lines show the 10^th^, 25^th^, 50^th^, 75^th^ and 90^th^ percentiles.

CV2 was calculated as 2 x |interval_n+1_ – interval_n_| / (interval_n+1_ + interval_n_).

## Acknowledgements

The authors wish to thank Dr. Ruben van der Giessen for fruitful discussions, Roberta Mazza for help with data analysis and Mandy Rutteman, Elize Haasdijk and Erika Goedknegt for technical support. Financial support was provided by the Netherlands Organization for Scientific Research (NWO-ALW; CIDZ), the Dutch Organization for Medical Sciences (ZonMW; CIDZ), Life Sciences (CIDZ), ERC-adv and ERC-POC (CIDZ).

## Author contributions

V.R., A.L.R., M.N., L.W.J.B. and C.I.D.Z. designed the paradigms, V.R., A.L.R., S.C. and L.W.J.B. performed the experiments, A.L.R. and M.N. provided new analysis tools, V.R., A.L.R. and S.C. performed the formal analysis, V.R., A.L.R., M.N., L.W.J.B. and C.I.D.Z. analyzed and interpreted the data, V.R., A.L.R., L.W.J.B. and C.I.D.Z. wrote the manuscript with input from all other authors, and C.I.D.Z. provided funding.

---

**Figure S1.**
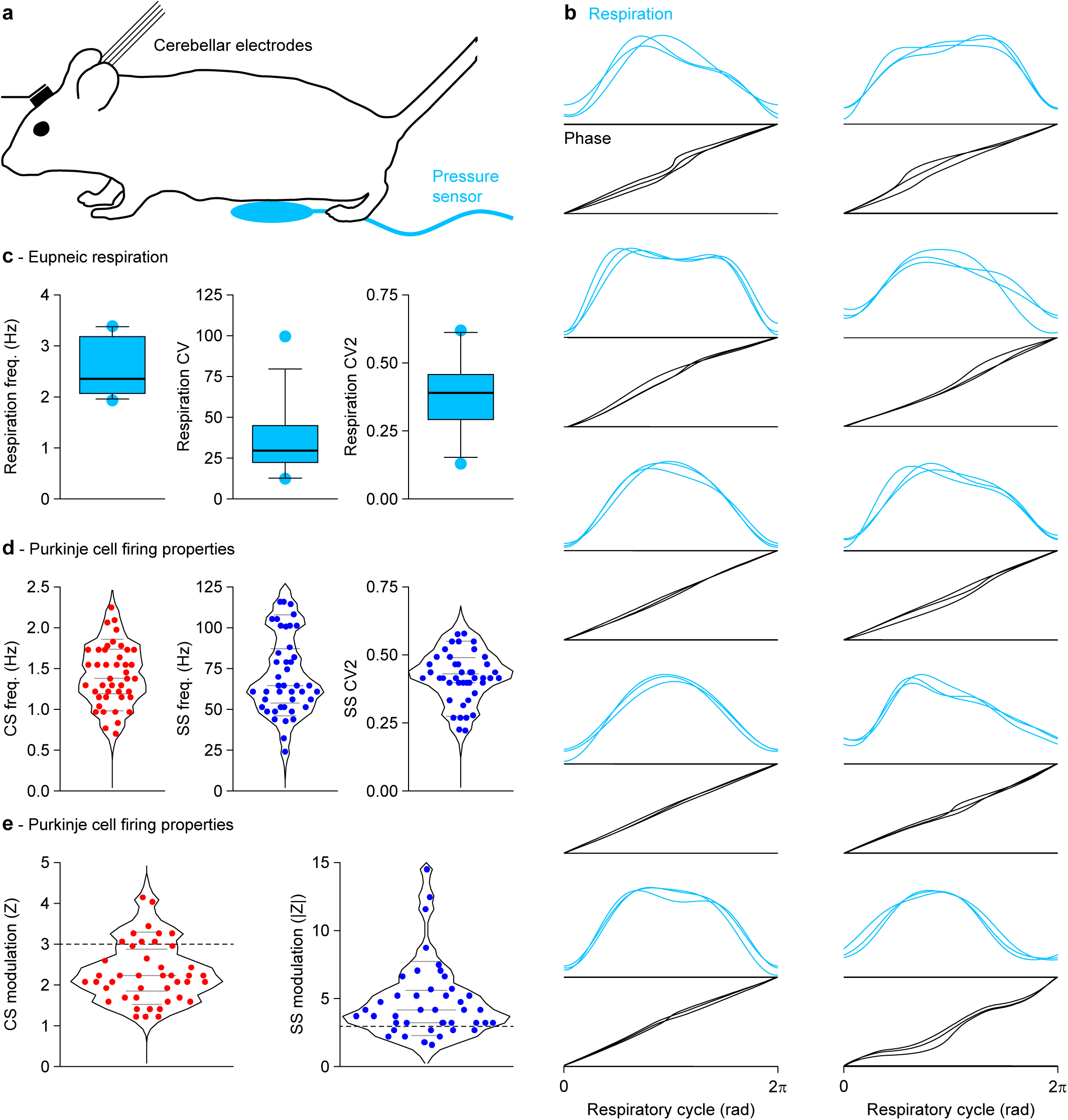
Purkinje cells in the lateral cerebellum encode eupneic breathing. **a** Single-unit recordings of Purkinje cells were made in the lobules simplex, crus 1 and 2 of awake, head-fixed mice during quiet, unperturbed (eupneic) respiration. **b** The pressure on the abdominal sensor was used as the raw respiratory signal (cyan). As the course and duration of each cycle could be quite variable, we used a phase transform (black) to obtain the instantaneous phase at each moment of the respiratory cycle. For ten mice, we show here three overlaid randomly selected cycles (during unperturbed breathing) with underneath it the three phase transforms. The three parts of the cycle, inspiration (starting at phase 0), post-inspiration and expiration can be seen in most traces. **c** Frequency, coefficient of variation (CV) and mean local coefficient of variation (CV2) of eupneic respiration in 13 mice. **d** The average complex spike (CS) and simple spike (SS) frequency as well as the mean local coefficient of variation (CV2) of the simple spikes of 43 Purkinje cells recorded during eupneic respiration. **e** Violin plots indicating the distributions of the maximal (absolute) complex spike and simple spike modulation for each Purkinje cell during the respiratory cycle. The firing rate modulation is expressed as Z score related to the bootstrap analysis. Responses exceeding a Z score of 3 (*p* < 0.01) were considered to be statistically significant, but it is clear that most Purkinje cells show at least some degree of modulation and any clear separation between modulating and non-modulating Purkinje cells would be subjective. Gray lines in the violin plots indicate 10^th^, 25^th^, 50^th^, 75^th^ and 90^th^ percentiles.

**Figure S2.**
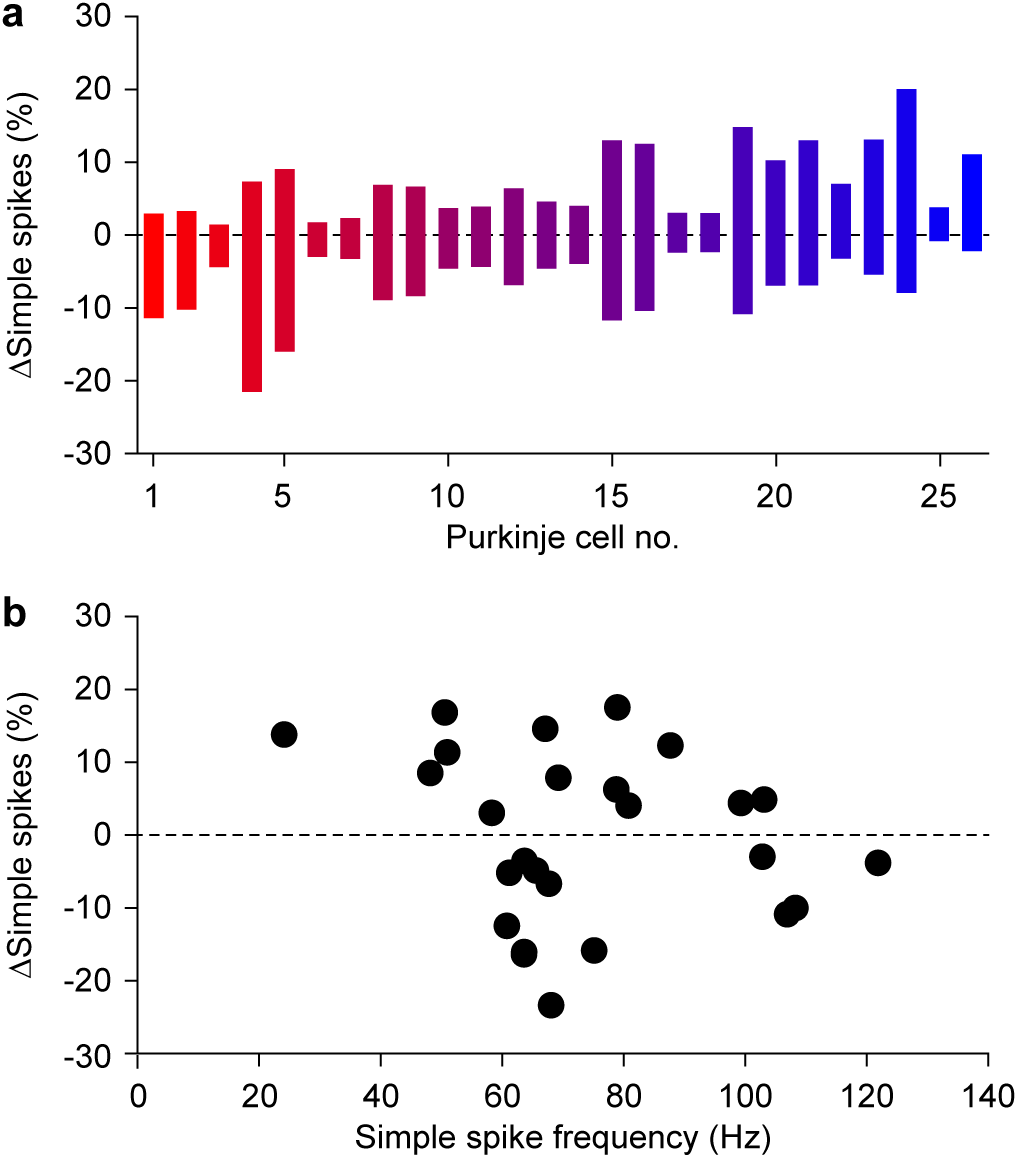
Eupneic respiration is associated with increased and decreased simple spike firing. **a** Maximal increase and decrease in simple spike firing in response to air puff stimulation per Purkinje cell. The cells are sorted based upon their bias towards decreased (left) of increased (right) simple spike firing. The Purkinje cells have the same color code as in Fig. 2c. **b** No correlation between average firing rate and maximal modulation in simple spikes (*p* = 0.309, Pearson correlation test).

**Figure S3.**
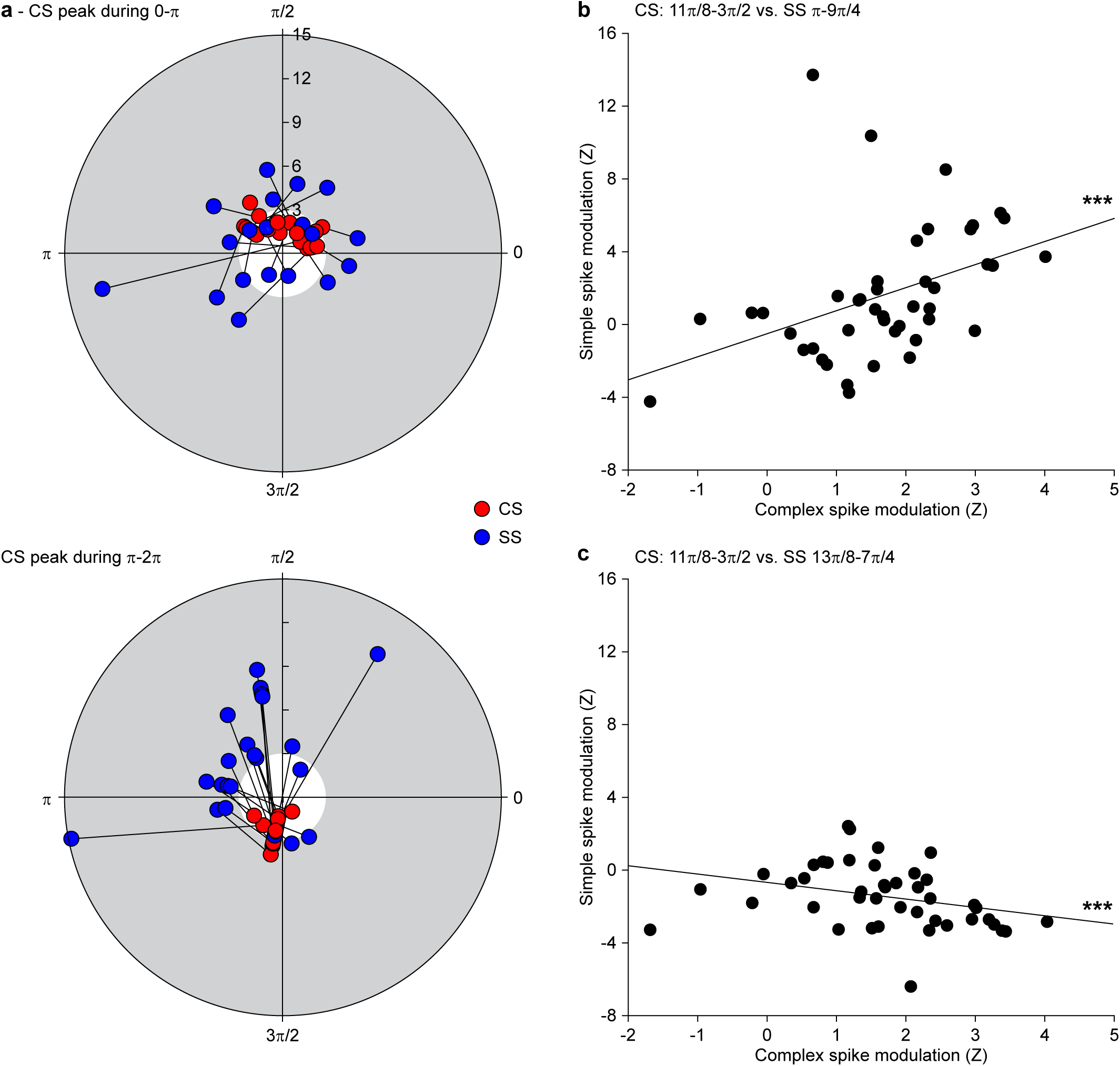
Complex spike and simple spike modulation occur during distint phases of the respiratory cycle. **a** Polar plot showing, for each Purkinje cell, the relation between the phase of maximal complex spike (red) and that of the strongest simple spike (blue) modulation during unperturbed respiration. The radial axis indicates the modulation strength (in absolute Z score). The grey area indicates |Z| > 3. The neurons are separated based on the occurrence of the peak complex spike modulation during the first (top) or second half (bottom) of the respiratory cycle. **b** There was a positive correlation between the rate of simple spike firing around the transition between inspiration and post-inspiration (∼π) and the occurrences of complex spikes during the transition from post-inspiration to expiration (∼3π/2) (r = 0.54, *p* < 0.001, Spearman rank test). **c** Likewise, there was a negative correlation between complex spike firing around the transition from post-inspiration to expiration and the simple spike rate during expiration (∼7π/4) (r = −0.43, *p* = 0.004, Spearman rank test).

**Figure S4.**
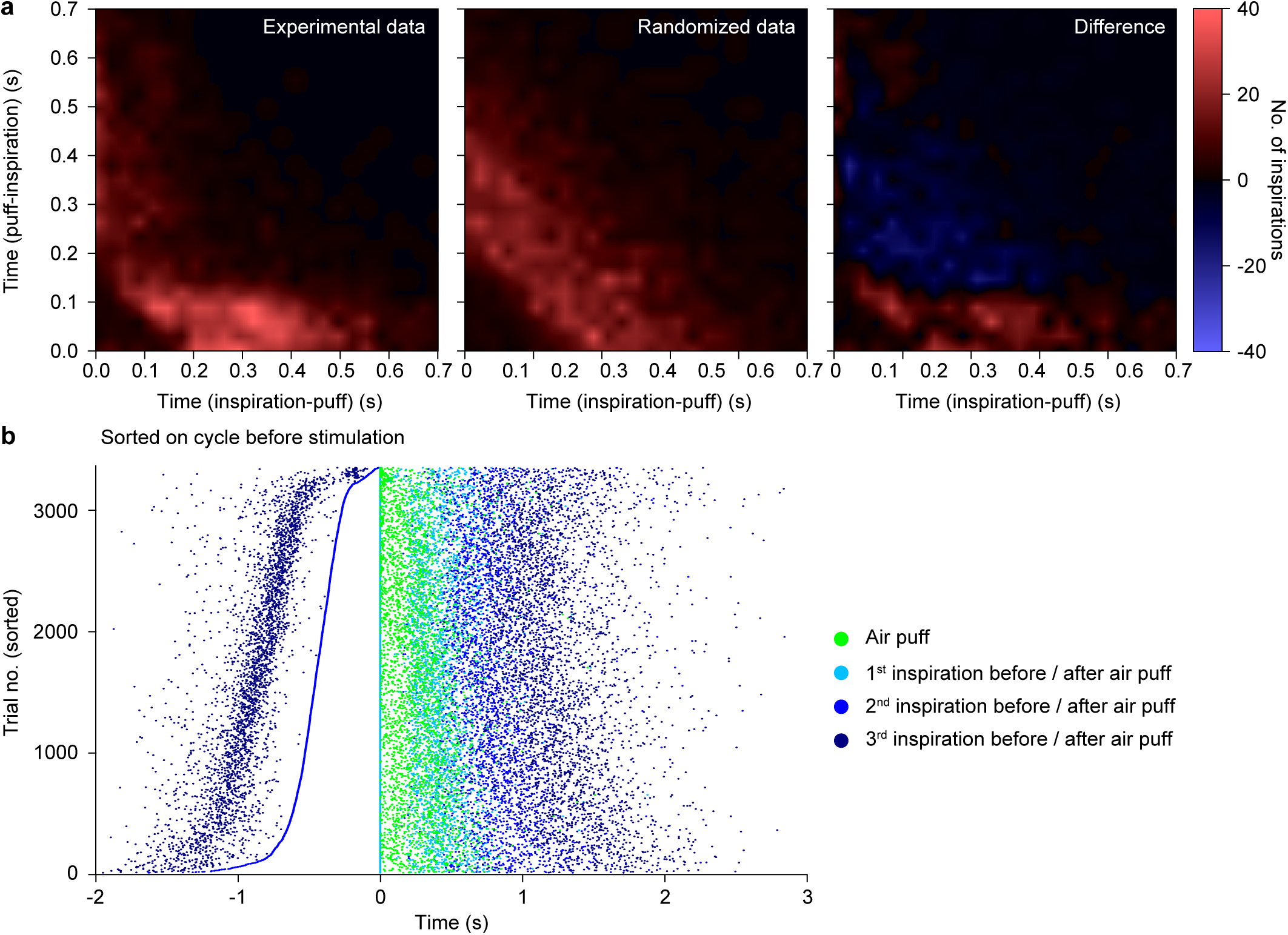
Whisker pad stimulation triggers inspiration. **a** Heat maps showing the distributions of the intervals between whisker pad stimulation and the start of the next inspiration (x axis) and the intervals between the start of the preceding inspiration and the whisker pad stimulation (y axis). The recorded data were compared to data were the times of the inspiration were randomly shuffled (cf. Fig. 4e-f). In the randomized data (middle), there is a clear symmetry between the time interval between the onset of inspiration and that of the stimulus (”inspiration - puff”) and the time interval from stimulus onset to the start of the next inspiration (”puff - inspiration”). This symmetry is broken in the experimental data (left), showing a tendency to start the next inspiration within 100 ms of the stimulus (right). **b** Whisker pad air puff stimulation affected the timing of the subsequent inhalations, but the mice did not entrain their respiration on the fixed frequency of the air puff stimulation. This becomes clear from the raster plot showing the timing of the start of inspiration around the moment of air puff stimulation. The raster plot is constructed by combining trials from 12 mice, sorted on the duration of the cycle prior to the air puff stimulation.

**Figure S5.**
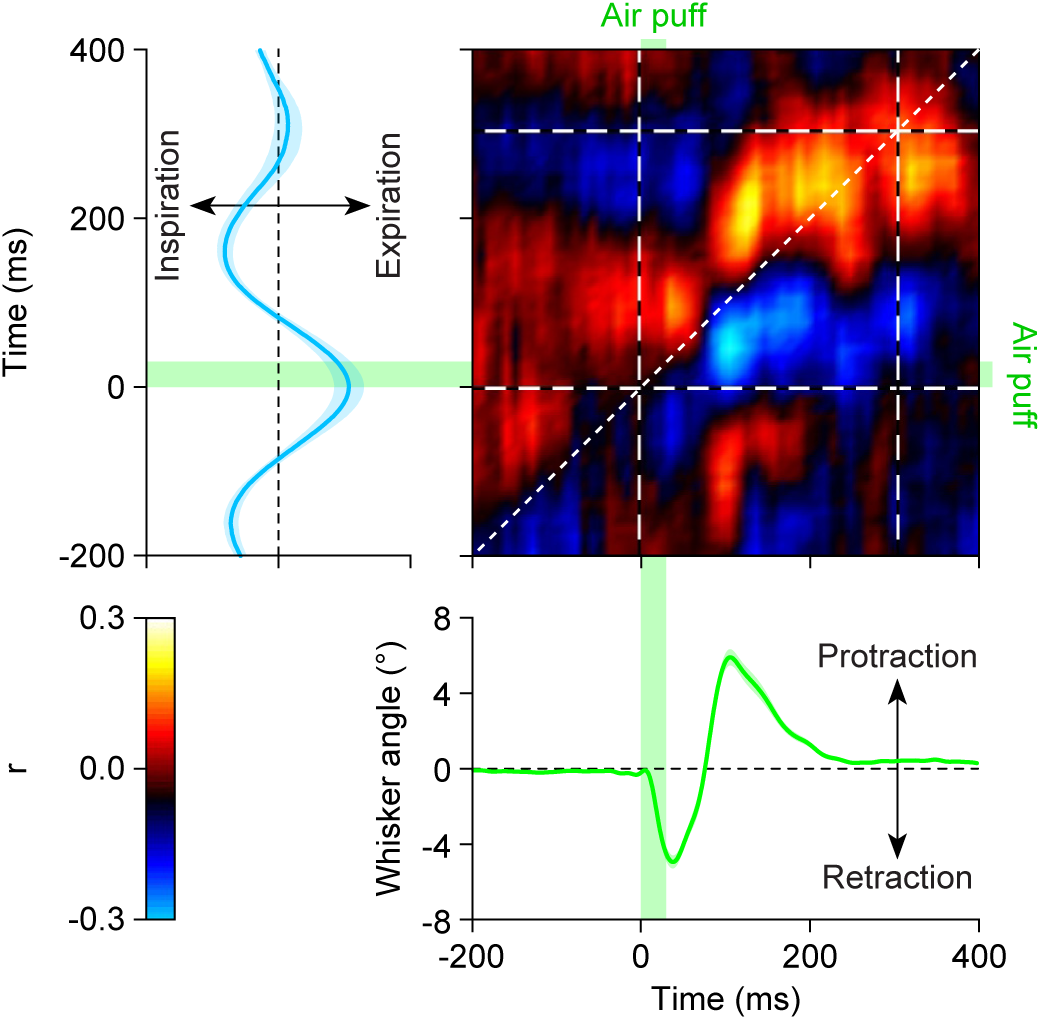
Accelerated inspiration follows reflexive whisker protraction. Air puff stimulation of the whikser pad induces a reflexive protraction of the facial whiskers that follows an initial, largely passive retraction (green trace, bottom). The same stimulus also accelerates inspiration (cyan trace, left). Trial-by-trial variance analysis indicates that the execution of both behaviors are correlated: whisker protraction is linked with a delay to inspiration. The heat map and the traces are the averages of the 11 mice for which whisker data, of 100 trials per mouse, were available (shaded areas: SEM).

**Figure S6.**
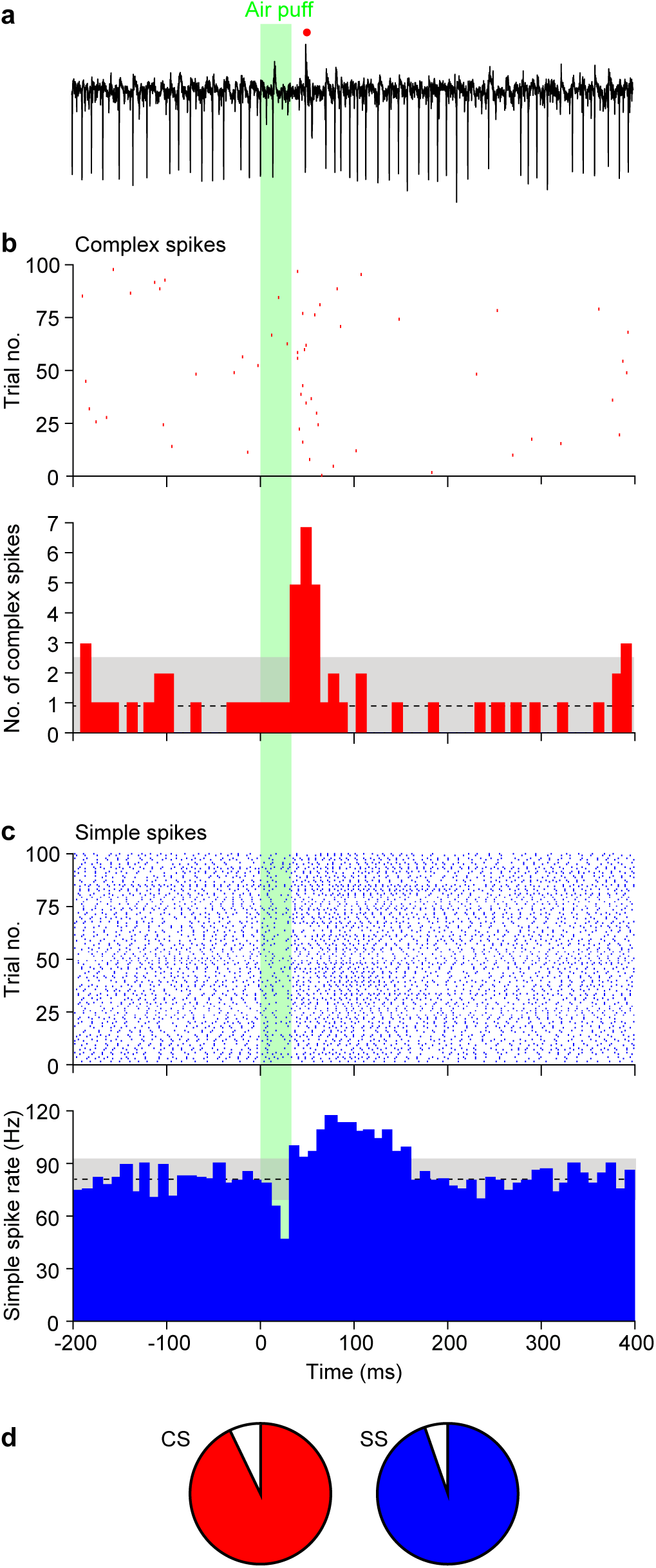
Purkinje cells respond to whisker pad air puff stimulation. **a** Extracellular recording of a representative Purkinje cell in crus 1 during air puff stimulation of the ipsilateral whisker pad. Of this same cell, raster plots and peri-stimulus time histograms of the complex spikes (**b**) and simple spikes (**c**) were made. Note the bidirectional modulation of the simple spikes. **d** Of the 57 recorded Purkinje cells, 53 (93%) responded with a statistically significant complex spike response to the whisker pad air puff. For the simple spikes, this number was 54 (95%).

**Figure S7.**
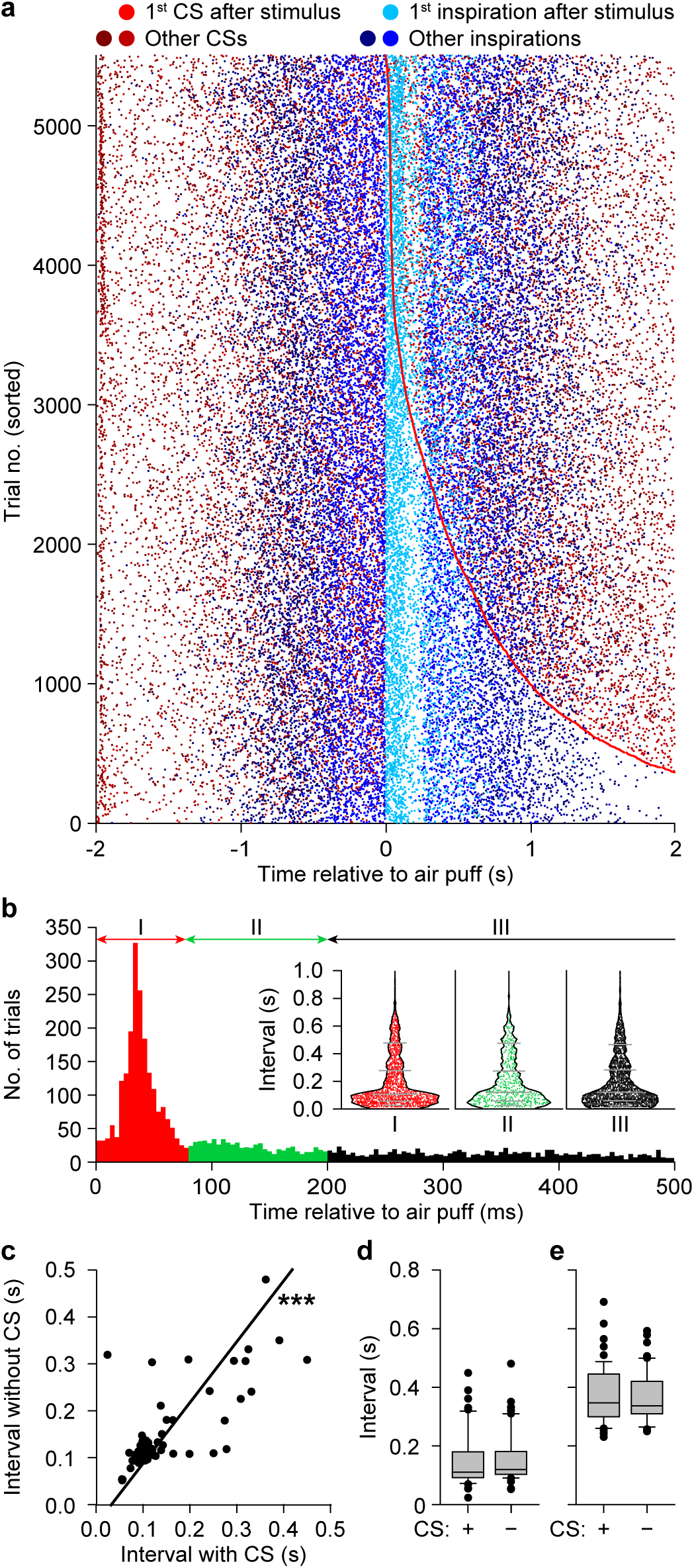
Complex spikes do not mediate the accelarated inspiration after whisker pad stimulation. **a** Air puff stimulation of the whisker pad triggers both complex spike firing (red dots) and accelerated inspiration (cyan dots). This raster plot shows the pooled trials of the 12 mice ordered on the interval between the start of the stimulus and the first complex spike afterwards. **b** Histogram of complex spikes during the first 500 ms after the air puff, composed of the data shown in **a**. The initial peak response occurs within 78 ms. Inset: Violin plots showing that the timing of the first inspiration after the air puff is not depending on the moment of complex spike firing. Left: trials with a complex spike between 0 and 78 ms after the air puff; middle: 78-200 ms; right: 200-500 ms. *p* = 0.560, KW = 1.158, Kruskal-Wallist test. **c** Scatter plot showing, for each Purkinje cell, the median interval between air puff and start of the next inspiration for trials with and without a complex spike within 78 ms of the air puff. The strong correlation demonstrates a lack of impact of complex spike firing on the start of the next inspiration (r = 0.693, *p* < 0.001, Spearman rank test). **d** Box plots of the intervals between the air puff and the start of the next inspiration in trials with and without a complex spike during the first 78 ms after the air puff (*p* = 0.148, Mann-Whitney test). **e** The same for the second respiratory interval after the air puff (*p* = 0.302, Mann-Whitney test).

**Figure S8.**
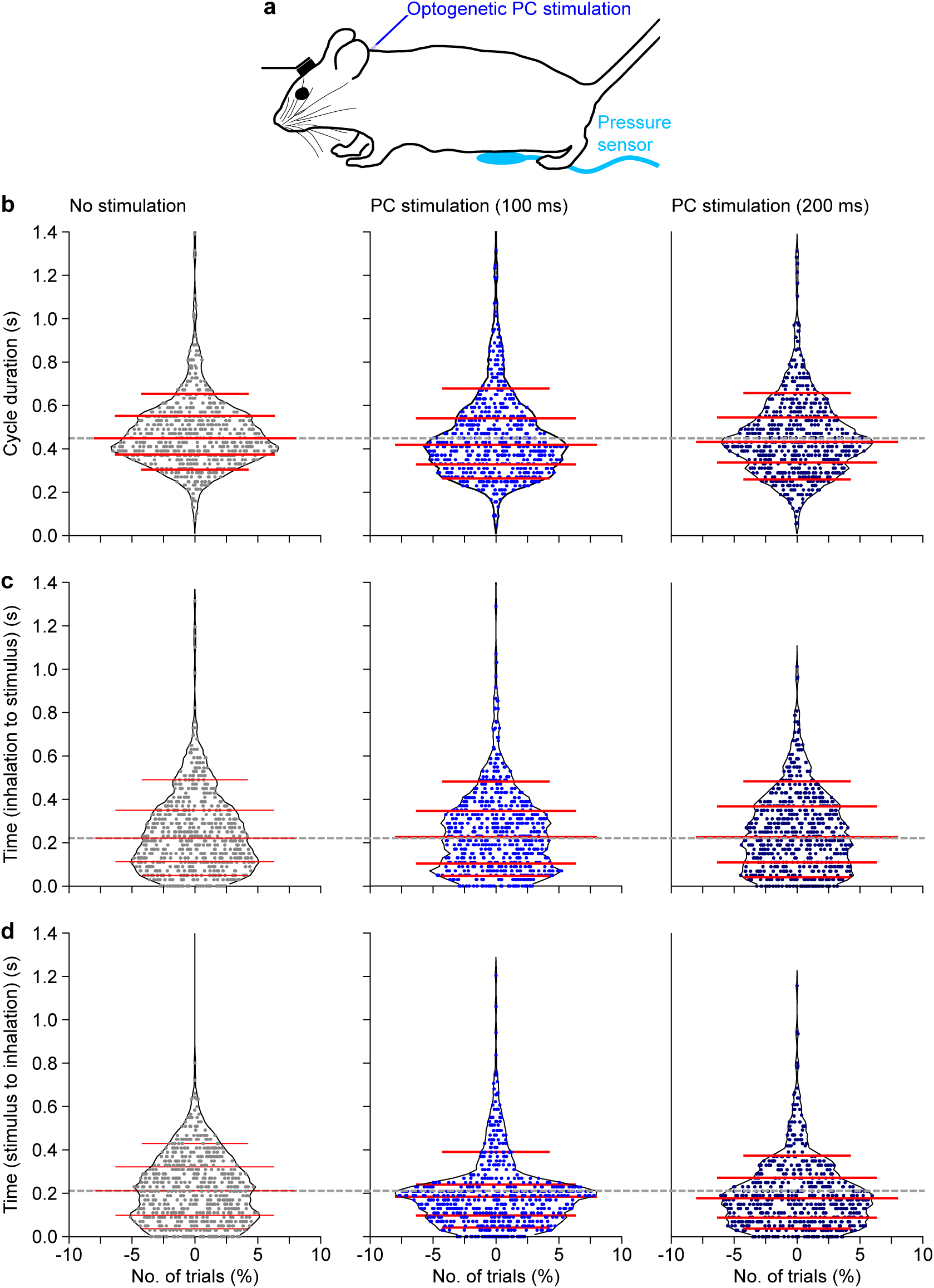
Purkinje cell stimulation affects respiratory timing. The impact of optogenetic stimulation on respiratory timing was studied using transgenic mice expressing ChR2 exclusively in their Purkinje cells (**a**). Violin plots showing the duration of the respiratory cycle during which the stimulus was given (**b**), the interval between the start of inspiration to that of the stimulus (**c**) and the interval between the start of stimulation and that of the next inspiration (**d**). Left column: 100 ms stimulation, right column: 200 ms stimulation. The horizontal lines indicate the 10^th^, 25^th^, 50^th^, 75^th^ and 90^th^ percentiles.

